# Neurophysiological markers of successful learning in healthy aging

**DOI:** 10.1101/2022.07.18.500426

**Authors:** Dawid Strzelczyk, Tzvetan Popov, Simon P. Kelly, Nicolas Langer

## Abstract

The capacity to learn and memorize is a key determinant for the quality of life, but is known to decline to varying degrees with age. Previous ERP research methods had the limitation that their design did not allow to track the gradual memory formation process. Thus, the neural mechanisms underlying memory formation and the critical features that determine the extent to which aging affects learning are still unknown. By using a visual sequence learning task, which consists of the repeated presentation of a simple sequence of tokens, we are able to track the progress of gradual memory formation through both neurophysiological and behavioral markers. On a neurophysiological level, we focused on two learning related centroparietal ERP components: the P300 and broad positivity.

Our results revealed that although both age groups showed significant learning progress, young individuals learned faster and remembered more stimuli than older participants. Successful learning was directly linked to a decrease of P300 amplitude. However, young participants showed larger P300 amplitude with a sharper decrease during the memory formation process. The P300 amplitude predicted learning success in both age groups, was associated with increased fronto-parietal brain network activation and showed good test-retest reliability. Highly, similar results were found for the broad positivity component, which raises the questions if the BP is a distinct component or just a prolonged P300. In a series of analyses, including topographic analysis of variance (TANOVA), equivalence testing and source reconstruction analysis, we addressed the unresolved questions. These analyses revealed concordant distributed brain activation patterns within parietal circuits. Thus, there is no evidence (rather evidence for equivalence) for distinct underlying neural generators for the two components.

Taken together, the results highlight the importance of the P300 as a neurophysiological marker of learning and may enable the development of preventive measures for age-related impeded learning.

## 1. Introduction

The aging of our population, with the increasing prevalence of physical and cognitive impairments, poses an increasing challenge to society. Aging is associated with decline in various domains including episodic memory and learning (DeCarli, 2003; Grady et al., 2010). However, there is a substantial variation in the rate, the onset, and the severity of the decline of learning and memory, ranging from mild (i.e., healthy aging) to dramatic cognitive impairments (e.g., Alzheimer’s disease (AD)).

In neuropsychology, learning capacity is mostly assessed by theoretically founded psychological test batteries. These tests have the advantage that the outcome measures typically hold high standards of psychometric properties (i.e., validity and reliability), and that individual outcome measures can be referenced to scores of a norm population. However, the performance measured by psychological tests usually reflects the product rather than processes of successful or impeded learning. As such, psychological tests generally provide little information about the processes of age-related decline in learning (Luu et al., 2009). Noninvasive neurophysiological methods, such as electroencephalography (EEG), may overcome these problems by providing insights into the mechanisms of successful or age-related impediments to learning and memory formation processes (Chiang et al., 2018; Tinga et al., 2019). EEG is particularly suitable for tracing the dynamics of memory formation processes at the fast pace typical of learning, owing to its high temporal resolution. Event-related potential studies have commonly found associations between memory formation and centro-parietal, positive potentials. Two key centro-parietal components are the P300 (here, we refer to the P3b as P300 in the remainder of this article) and broad positivity (BP).

The P300 component is a centro-parietal positivity peaking around 300 ms post-stimulus onset (Sutton et al., 1965). The P300 is elicited only by task-relevant stimuli and its amplitude decreases as a function of the prior probability of a stimulus, thus providing a neural index of stimulus expectancy (Donchin, 1981; Duncan-Johnson & Donchin, 1977; Kolossa et al., 2012; Mars et al., 2008; Rüsseler et al., 2003; Steinemann et al., 2016). In the context of sequence learning, expectancy grows alongside the accumulation of knowledge about the learned material: known stimuli are more expected than unknown stimuli. Although several studies have reported an age-related increase of the P300 latency with a concurrent decrease of its amplitude in adults (Fjell & Walhovd, 2001; Polich, 1997, 2007; Porcaro et al., 2019; Tinga et al., 2019; van Deursen et al., 2009), it is not yet known how learning-related modulations of the P300 change with age.

In addition, early memory-related ERP studies reported larger broad centro-parietal positivity (BP) peaking between 300 and 800 ms post-stimulus onset in young individuals for words that were later remembered compared to those later forgotten (Karis et al., 1984; Neville et al., 1986; Paller et al., 1987). Latency and amplitude of this component varied depending on the paradigm and stimulus properties (Johnson, 1995). Later, (Steinemann et al., 2016) demonstrated that the BP amplitude was especially elevated for newly learned stimuli. And yet, the question whether BP is a distinct component or just a prolonged P300 is still open to debate (Hajcak & Foti, 2020; Johnson, 1995; Verleger, 2020)

Previous memory formation research on the P300 and BP was conducted mainly in young individuals. Consequently, it is largely unknown how aging affects the modulation of P300 or BP amplitudes and latencies over the course of learning. Therefore, the contribution of the present study is to investigate age-effects of these neurophysiological measures related to learning. By using a visual sequence learning task, which consists of the repeated presentation of a simple sequence of tokens, we are able to track the progress of gradual memory formation through both neurophysiological and behavioral markers. First, we expect both behavioral learning performance, defined as the cumulative sequence knowledge, and learning rate to be better in young compared to older individuals. We further predict that these behavioral indices are directly related to neurophysiological characteristics of the ERP components (i.e., P300 and BP amplitudes). More specifically, the sequence knowledge can relate to correct stimulus expectancy. Hence, we hypothesize that the P300 amplitude will decrease monotonically over the course of learning, from stimuli being unknown, to newly learned and finally to become fully known. On the other hand, we predict the BP amplitude, hypothesized here as the signal of active memory trace formation, to be especially elevated for stimuli currently being committed to memory (i.e., newly learned). Overall, we predict similar patterns of neural activation in both age groups, but decreased amplitudes and increased latencies in older individuals compared to the young individuals. Further, we examine if P300 and BP are distinct components or just a prolonged P300 by using equivalence tests on sensor level and the comparison of the source reconstruction of the two components. Finally, we examine the potential of the neurophysiological measures to predict learning success within and across individuals. Such prediction from neurophysiological measures, independent of behavioral indices, could facilitate the development of new diagnostic tools for age-related learning difficulties.

## 2. Methods

### 2. 1. Participants

In the present study 100 young and 117 older participants were recruited. Data from one young and two older participants were excluded from further analysis due to technical problems (see exclusion criteria below). This resulted in a remaining sample of 99 young (age range 19.7 - 43.2 years; mean age, 24.97 ± 4.72 years, 39 male, 77 right-handed) and 115 older individuals (58 - 83.5 years; mean age, 69.1 ± 5.35 years, 55 male, 98 right-handed). Table 1 shows basic demographic details for both age groups. All participants were healthy, reported normal or corrected to normal vision and no current neurological or psychiatric diagnosis. The young group consisted of graduate students at the University of Zürich or other universities nearby. The older subjects were recruited during lectures within the Senior-University of Zürich. As a compensation the participants were given course credit or monetary reward (25 CHF/h). This study was conducted according to the principles expressed in the Declaration of Helsinki. The study was approved by the Institutional Review Board of Canton Zurich (BASEC - Nr. 2017 - 00226). All participants gave their written informed consent before participation.

**Table 1.** Demographics information

|  | *Young* |  | *Older* |  | *t* | *df* | *p-value* |
| --- | --- | --- | --- | --- | --- | --- | --- |
|  | *Mean* | *SD* | *Mean* | *SD* |  |  |  |
| Gender | 39m/62f | | 55m/64f | | $\chi^2=1.29$ | | 0.256 |
| Age (years) | 24.97 | 4.72 | 69.04 | 5.35 | -63.42 | 212 | 8e-140*** |
| Years of Education | 15.11 | 3.2 | 15.39 | 3.29 | -0.64 | 210 | 0.521 |
| MMSE |  |  | 29.02 | 1.02 |  |  |  |

\* $p < 0.05$ . \*\* $p < 0.01$ . \*\*\* $p < 0.001$

### 2. 2. Procedure

The data used in this study was recorded in our laboratory in the context of a larger project. This larger research project aims to quantify age effects on eye movement behavior and electroencephalography (EEG) recordings of resting-state and other task-based paradigms. In addition, the reliability of behavioral and neurophysiological measures is also assessed. For this reason, the data was collected in two experimental sessions separated by a week. Older participants were given a questionnaire to complete at home regarding basic demographics, handedness, social interactions, and social status. Upon arrival, each participant’s cognitive functioning was tested with a battery of neuropsychological tests that lasted for about 35-45 minutes. Additionally, the older participants performed the Mini-mental State Exam (MMSE) in order to screen for cognitive impairment and dementia (Folstein et al., 1975). All participants accomplished a MMSE score above the threshold of 25.

Subsequently, the participants were comfortably seated in a chair in a sound- and electrically shielded Faraday recording cage. The cage was equipped with a chinrest to minimize head movements and a 24-inch monitor (ASUS ROG, Swift PG248Q, display dimensions 531 x 299 mm, resolution 800 x 600 pixels resulting in a display: 400 x 298.9 mm, vertical refresh rate of 100 Hz) on which the experiment was presented. The distance between the chinrest and the monitor was 68 cm.

### 2. 3. Sequence Learning Task

An explicit visual sequence learning paradigm was first developed by (Moisello et al., 2013) and is currently considered as an important tool in assessing reliable indices of memory formation and learning progress (Steinemann et al., 2016). The advantage of this paradigm is the simplicity of the stimuli, which enable it to differentiate cortical computations associated with perceptual memorization and stimulus identification (Langer et al., 2017). The participants were asked to learn a fixed sequence of eight visual stimulus positions (Figure 1A). The stimuli consisted of filled white circles (visual angle of 0.84°) and were presented on a computer screen with a bright gray background, positioned equidistant around a ring of fixed eccentricity (visual angle of the distance between center of the screen and the stimulus of 4.21°). Each stimulus was presented for 600 ms with an offset-to-onset interval of 1300 ms. Before the main task recording, a training task was administered, consisting of 4 stimuli placed on the same 8 locations, in order to familiarize the participants with the tasks and to ensure task comprehension. The participants performed the training task until they correctly recalled all 4 locations. Feedback was provided only during the training task.

**Figure 1.**
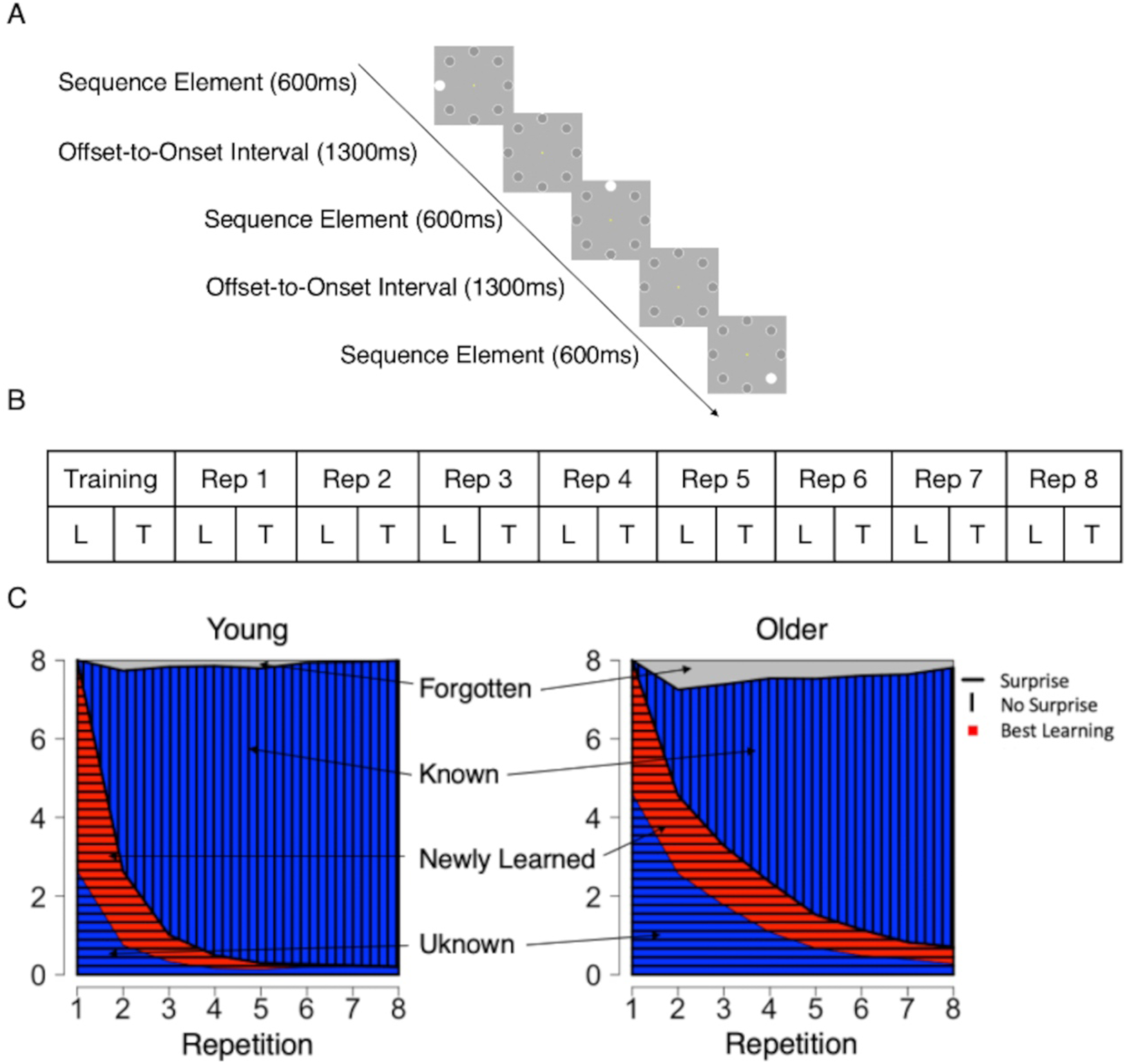
Sequence learning task and the design of the present study. (A) A sequence consisted of eight positions at which white circles were presented one by one. Each stimulus was presented for 600 ms with the interstimulus interval of 1300 ms. (B) Each sequence repetition (Rep) consisted of a learning phase (L) in which all sequence elements were presented and a testing phase (T) in which the participants were asked to recall the position of the stimuli. (C) Development of the average number of stimuli in each learning category over sequence repetitions for young (left) and older (right) individuals. Average was taken across participants and all six stimulus sequences. Grating indicates learning states in which the participants were surprised about the presented stimuli (i.e., unknown, newly learned; horizontal grating) or had relatively accurate knowledge (i.e., known; vertical grating). The red color indicates the process of committing the stimulus to memory (i.e., newly learned).

The main task consisted either of eight sequence repetitions or ended after the participant correctly recalled the sequence of stimuli three times in a row. Each sequence repetition consisted of a learning phase and a testing phase. In the learning phase, the participants were told to focus on a yellow dot at the center of the screen (controlled by an eye-tracking device) and memorize the position of each stimulus. In the test phase, the participants attempted to recall the sequence using a computer mouse by clicking the locations on a computer screen. There was no time restriction on providing responses, and no feedback on their performance was given. The duration of the paradigm for learning a sequence varied between 2 and 5 min, depending on the speed of recall reports and number of sequence repetitions (i.e., 3 to 8 repetitions). Overall, each participant learned six different sequences, resulting in a minimum of 18 (in case the participant solves everything correctly from the beginning) and maximum of 48 sequence repetitions.

### 2. 4. Behavioral Data

Each individual stimulus presentation was classified as one of four categories based on the responses of the participants after each sequence repetition. A stimulus was assigned to the *unknown* category (UN) when location was recalled incorrectly in the current and all previous repetitions, to the *newly learned* category (NL) when stimulus position was recalled correctly for the first time in the current repetition, to the *known* category (K) when stimulus position was correctly recalled at least twice in a row and to the *forgotten* category (F) when stimulus position was once recalled correctly but incorrectly in the following sequence repetition. The *forgotten* category was excluded from further analysis due to insufficient number of trials (on average 4.7% per participant).

Two behavioral performance indices were computed for each participant. The *knowledge index* (KI), which reflects the cumulative sequence knowledge, was defined as the ratio of the number of known nK(P, sr) and newly learned stimuli nNL(P, sr) to the total number of stimuli nT(P, sr) in each sequence repetition, where P represents the participant and sr the sequence repetition. The newly learned stimuli were weighted by 0.5 to take into account the fact that they might be classified as known 50% of the time during the transition from unknown to known.

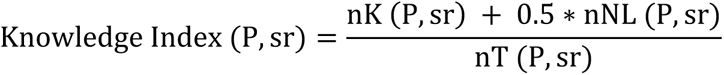

The *learning index* (LI), which reflects the learning rate in each sequence repetition, was defined as the ratio of the number of newly learned stimuli to the total number of stimuli in each sequence repetition.

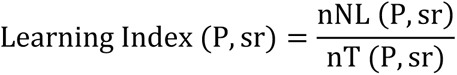

#### 2. 5. 1. EEG data acquisition

EEG data were recorded at a sampling rate of 500Hz using a 128-chanel Hydrogel net system (Electrical Geodesics Inc.). The recording reference was at Cz (vertex of the head), and impedances were kept below 40 kΩ.

#### 2. 5. 2. Eye-Tracking data acquisition

An infrared video-based eye tracker (EyeLink 1000 Plus, SR Research; http://www.srresearch.com/) recorded eye movements at a sampling rate of 500 Hz and an instrumental spatial resolution of 0.01°. The eye tracker was calibrated and validated before each task with a 9-point grid until the average error for all 9 points was below 1°. In this study eye tracker data were used in a control analysis accounting for whether the participants maintained focus at the center of the screen as instructed (i.e., yellow dot). Therefore, eye movements during each stimulus presentation period were included into the models as a covariate (i.e., 1 = kept fixation, 0 = lost fixation). It was deemed that a participant had kept fixation, if the gaze was directed at the center of the screen within a square of side 1.26° of visual angle during at least 90% of the total duration of the stimulus presentation period.

#### 2. 5. 3. EEG data preprocessing

All data were analyzed using Matlab 2020a (The MathWorks, Inc., Natick, Massachusetts, United States) and RStudio 4.0.2 (R Core Team). The data were preprocessed in Automagic 2.4.3, a MATLAB based toolbox for automated, reliable and objective preprocessing of EEG- datasets (Pedroni et al., 2019). In the first step in Automagic, the bad channels were detected using the PREP pipeline (Bigdely-Shamlo et al., 2015). A channel was define as bad based on 1) extreme amplitudes (z-score cutoff for robust channel deviation of more than 5), 2) lack of correlation (at least 0.4) with other channels with a window size of 1s to compute the correlation, 3) lack of predictability by other channels (channel is bad if the prediction falls below the absolute correlation of 0.75 in a fraction of 0.4 windows of a duration of 5s), 4) unusual high frequency noise using a z-score cutoff for SNR of 5. These channels were removed from the original EEG data. The data was filtered using a high-pass filter with 0.05 Hz cutoff and a low-pass filter with 45 Hz cutoff using the EEGLAB function pop_eegfiltnew (Widmann & Schröger, 2012). The high-pass filter was chosen based on (Tanner et al., 2015) that recommended a range of 0.01-0.1 Hz for slower cortical potentials such as the P300. Line noise was removed using a ZapLine filter with a passband edge of 50 Hz (de Cheveigné, 2020). Next, independent component analysis (ICA) was performed. However, as the ICA is biased towards high amplitude and low frequency noise (i.e., sweating), the data was temporarily filtered with a high-pass filter of 1 Hz in order to improve the ICA decomposition. Using the pre-trained classifier IClabel (Pion-Tonachini et al., 2019) each independent component with a probability rating >0.8 of being an artifact such as muscle activity, heart artifacts, eye activity, line noise and channel noise were removed from the data. The remaining components were back-projected on the original 0.05 Hz high-pass filtered data. In the next step, the channels identified as bad were interpolated using the spherical interpolation method. Finally, the quality of the data was automatically and objectively assessed in Automagic, thus increasing research reproducibility by having objective measures for data quality. Using a selection of 4 quality measures the data was classified into three categories: Good (1070 sequences), OK (111 sequences) or Bad (24 sequences). Data was classified as bad, if 1) the proportion of high-amplitude data points (>30μV) in the signal is greater than 0.2, or 2) more than 20% of time points show a variance greater than 15μV across all channels, or 3) 30% of the channels show variance greater than 10μV, or 4) the ratio of bad channels is greater than 0.3. For the further analysis only the datasets with Good and OK ratings were used. Due to technical difficulties (EEG data not saved properly, missing participant’s responses, Matlab crash during experiment) data from 45 sequences was not available, resulting in a total of 1205 sequences (i.e., 47’213 stimuli).

Subsequent analyses were conducted using EEGLab (Delorme & Makeig, 2004), an open-source software for processing of electrophysiological signals. First, 23 channels were excluded from further analysis, including 10 EOG channels and 13 channels located on the chin and neck as they capture a little brain activity and are mostly contaminated with muscle artifacts (Langer et al., 2017; Tröndle et al., 2020, 2021). Next, the data was re-referenced to average reference and segmented from -100 to 800 ms after stimulus onset (i.e., presentation of white circle on one of the eight positions). The segments were inspected using an amplitude threshold of 90μV resulting in 5.59 % rejected segments on average for each participant.

As for baseline correction, rather than assume no systematic differences between stimulus categories and age groups in the baseline interval, the baseline was taken as a predictor into the linear mixed effect model, allowing the amount of baseline correction to vary across conditions (Alday, 2019). Otherwise, if there were systemically differences in the baseline interval between conditions, the traditional baseline correction would transfer these differences on later components. This approach has been proven superior outlined in a recent publication (Alday, 2019). The baseline was computed for each trial as an average of a centro-parietal electrode cluster in a time window of -100 to 0 ms.

#### 2. 5. 4. P300 and Broad Positivity extraction

Electrode sites and time windows for computing P300 and BP amplitude were extracted in the following analysis steps: First, to identify the electrode cluster for subsequent statistical analysis, which is representative for all participants, the grand average scalp topographies of unknown and newly learned trials between 200 ms and 800 ms with a 50ms step were plotted for young and older participants separately (Supplementary Figure 1A-B). Only the unknown and newly learned trials were selected, because the known trials were not expected to produce a strong P300 peak. Next, the maximal centro-parietal positivity (i.e., P300; ∼350 ms after stimulus onset) was identified and six electrodes (E54, E55, E61, E62, E78, E79) located around this centro-parietal positivity were chosen for computing the grand average ERP waveforms (Figure 3A). The electrode cluster was identical for both age groups.

**Figure 3.**
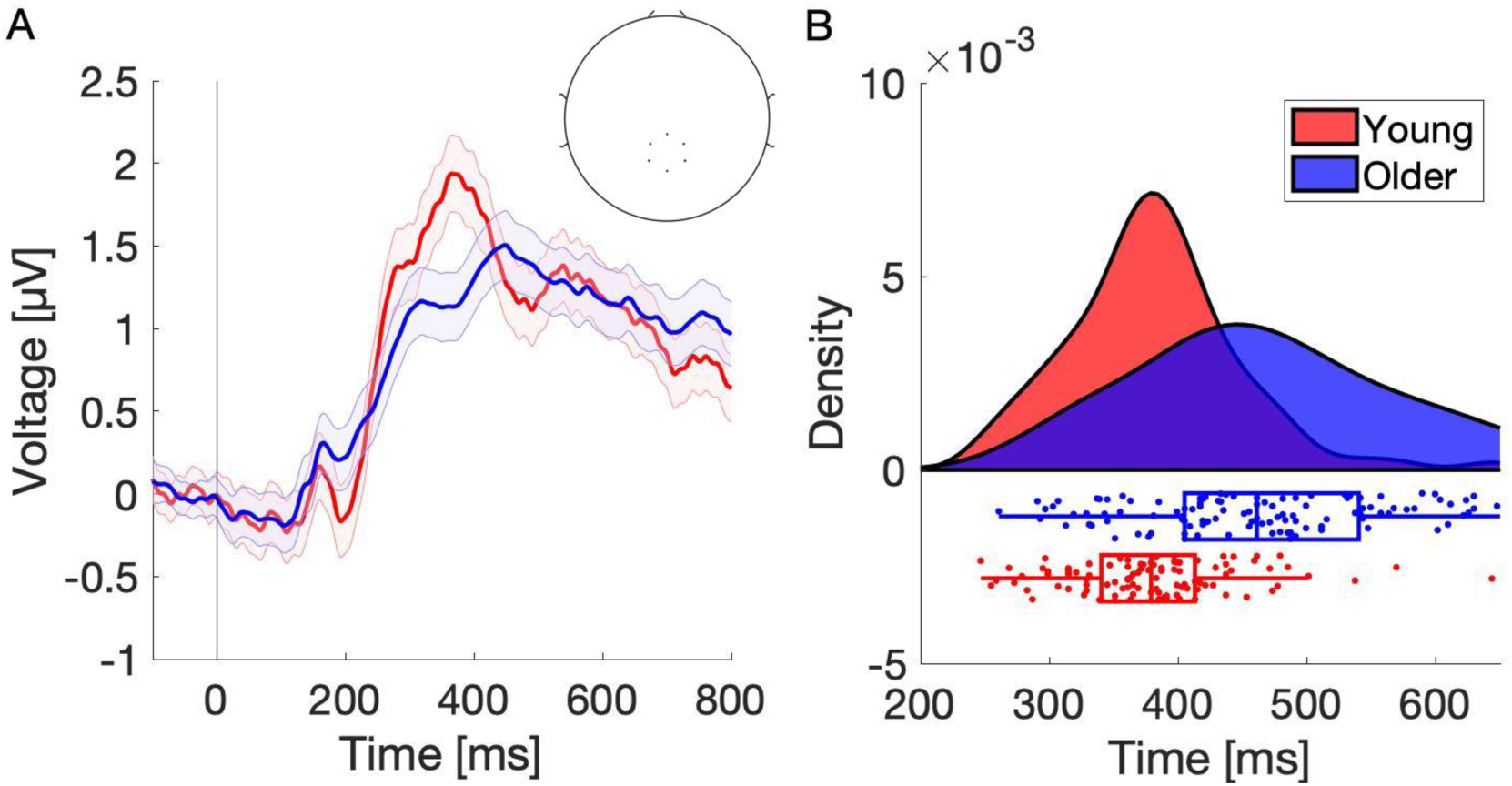
Grand average ERP and interindividual variability of P300 peaks in young (red) and older (blue) participants. (A) Grand average of young and older participants was computed using six centro-parietal channels: E54, E55, E61, E62, E78, E79 (see topoplot). Only unknown and newly learned trials were used, as the known trials were not expected to elicit a clear P300 peak. Shaded error bars represent the standard error of the mean. (B) Distribution of individual P300 peaks in young and older participants. Each dot on a boxplot represents a P300 50%-area peak latency. On average, young showed earlier P300 peak latency and smaller P300 peak variability (M = 373.1 ms, SD = 54.2 ms) compared to older participants (M = 459.9 ms, SD = 100.1 ms). The peak latencies were computed using the Liesefeld (2018) method based on unknown and newly learned trials. Please note that the age-related differences in amplitude visible in the grand average might be biased by higher P300 peak variability in older participants. To account for this, participant specific P300 amplitudes were calculated for all subsequent analyses.

However, it is well known that the P300 peak latency varies across individuals based on several factors such as age, gender, intelligence, memory capacity, personality and cognitive impairments including depression and dementia (Hansenne, 2000; Patterson et al., 1988; Polich, 2007). Furthermore, the intersubject variability of the P300 is somewhat enhanced in older subjects, which might reflect the larger variation of cognitive function in older individuals (Polich et al., 1985; Walhovd et al., 2008). The reported age-related increased variability of P300 peak latencies might introduce a bias and underestimate the true P300 amplitude in older individuals, when using a fixed time window for analysis, which is currently the common method in P300 research. For this reason, instead of using fixed time windows for statistical analysis based on grand average ERP, we computed an individual peak and time window for each participant using a method introduced by (Liesefeld, 2018)). The initial search window for individual peaks was defined based on the grand average peaks for each age group individually (P300: 250-490 ms in young, 320-560 ms in older; BP: 500-790 ms in young, 560-790 ms in older). From the Liesfeld approach, we extracted the 50%-area latency, onset and offset of a component for each participant. The 50%-area latency is the time point, in which a component has reached 50% of its area under the curve. The onset and offset of a component were defined by the time point where the component has reached 60% (and has fallen back below 60%) of its amplitude relative to pre-stimulus baseline (see Liesefeld for details). The 50%-area latency was then used as a measure of components latency and for the computation of the interindividual P300 peak latency variability. The individual onset and offset of a component were used to compute the average P300 and BP amplitudes for each participant’s trial (i.e., average over six centro-parietal electrodes between onset and offset of a component).

### 2. 6. Statistical analysis

Except for the interindividual variability of P300 peaks, for which we computed a Levene’s test (Olkin, 1960), we analyzed all behavioral and neurophysiological data using linear mixed effect models. The models were as general as possible at first and were progressively simplified using the step function for linear mixed effect models in R Studio (Kuznetsova et al., 2017) in order to identify the best-fit model. The predictors included the repetition number (continuous variable: 1-8), age group (factor of 2 levels: young, older), learning categories (factor of 3 levels: unknown, newly learned, known), baseline (continuous variable), session (factor of 2 levels: 1, 2), eye movements (factor of 2 levels: kept, lost fixation), gender (factor of 2 levels: male, female), stimulus number (factor of 8 levels, i.e., the presentation order of stimuli in a given repetition), subject (factor of 214 levels), sequence number (factor of 6 levels: 1-6, i.e., each participant attempted to memorize six sequences). The fixed and random effects of the model were specified depending on the goal of the analysis, having in mind that mixed model requires at least 5 levels for a random intercept term to accurately estimate the variance (Harrison et al., 2018). To overcome the increasing complexity of the models (due to inclusion of baseline) only the variables of interest and their interactions were further interpreted (i.e., repetition number, age group, learning states) and the covariates were included as controls. Furthermore, we did not examine the interaction effects of covariates such as gender, session and eye movements, because they are not in the main interest of this paper.

#### 2. 6. 1. Behavioral analysis

We began the behavioral analysis by computing the average number of stimuli per subject across learning categories (i.e., unknown, newly learned, known, forgotten) and age groups. Next, we examined the behavioral learning progress of both age groups. Learning performance was formally modeled as the cumulative knowledge about the sequence (i.e., knowledge index: KI) and the learning rate in each sequence repetition (i.e., learning index: LI). We specified two linear mixed effect models with a KI and LI as dependent variables (continuous variable: 0-1). The fixed effects included the sequence repetition number, age group, interaction of sequence repetition number and age group, session, gender and the random effects included subject and sequence number. In the following formulas fixed effects are denoted by a “+” symbol and interaction effects by an “∗” symbol, in line with the Wilkinson notation (Wilkinson & Rogers, 1973).

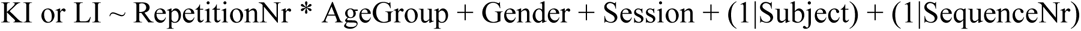

#### 2. 6. 2. EEG data analysis

##### Interindividual variability of P300 peak latency

To investigate age effects of the variability of interindividual peak latencies, we tested the equality of variances of both age groups by conducting the Levene’s test (Olkin, 1960).

##### P300 and BP amplitude over sequence repetitions

In the next step, assuming the inverse effect of expectancy on P300 amplitude, we investigated the changes in P300 and BP amplitude over sequence repetitions to test the hypothesis that P300 and BP decrease over the course of learning as the expectancy increases, and to assess their eligibility for prediction of learning success. First, we computed average P300 (i.e., mP300) and BP (i.e., mBP) amplitudes in each of the eight sequence repetitions. Subsequently, we defined a linear mixed model for both, average P300 and BP amplitude as dependent variables (continuous variable) and included repetition number, age group, average baseline in a given repetition, interaction of repetition number, age group and baseline, and additionally gender and session as fixed effects and the subject and sequence number as a random effect.

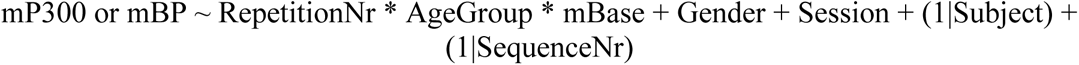

##### EEG signatures of successful learning

Next, we aimed to investigate whether the P300 and BP amplitudes are directly linked to the behavioral indices and could predict learning success and learning rate over sequence repetitions, respectively. P300 is modulated by stimulus expectancy and in sequence learning expectancy corresponds to increasing knowledge. Hence, we examined the relationship of the average P300 amplitude in each sequence repetition with the knowledge index. Meanwhile, the BP, as a signal associated with active memory trace formation, was expected to be especially elevated in repetitions where most stimuli were successfully committed to the memory; Hence we tested the relationship between BP and learning index. Using linear mixed models, we tested whether the KI and LI in a given sequence repetition can be predicted by the average P300 and BP in this repetition, respectively. Additionally, we used the age group, average baseline, gender and session as fixed effects and the subject and sequence number as random effects.

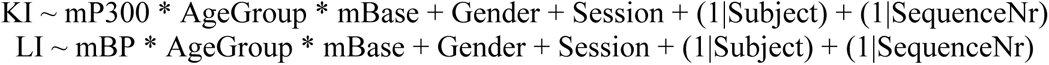

##### Predicting learning success across participants

For the prediction of learning success across participants, we fitted an exponential function to the learning curve, i.e., knowledge index, of each participant. Next, we obtained the time constant tau, which reflects the point at which the participant has learned the 63% or 1 – 1/e of the total sequence (Steinemann et al. (2016). Small tau values indicate fast learners. Also, other criterion values than 63% for defining tau were tested, such as 30%, 50%, 70%, in order to control the stability of this constant and their relation to the P300 amplitude. Next, the decrease in the P300 amplitude between the first and the third sequence repetition was computed, where most of the learning took place. Finally, we specified a linear mixed effects model with the time constant tau as the dependent variable (continuous variable). The fixed effects factors included the difference in P300 amplitude between the first and the third sequence repetition (continuous variable), age group, gender, session, and interaction of P300 difference, age group as fixed effects and the subject and sequence number as random effects.

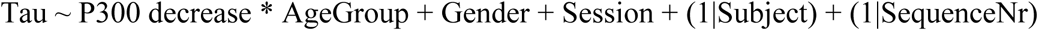

##### P300 and BP amplitude across learning categories and age groups

The decrease of P300 and BP amplitudes over sequence repetitions might be caused by habituation and occur purely as a function of time spent on the task. To provide stronger evidence for the relationship of both ERP components with learning, we examined the differences in P300 and BP amplitude across learning categories (i.e., unknown, newly learned, known stimuli). For this analysis, we conducted the linear mixed model on single trial data, because the trial-averaged ERP (e.g., averaging across trials within the same condition) has at least two limitations: first, the averaging approach has the drawback of different numbers of stimuli for each participant in each category, which has an impact on the signal-to-noise ratio. Second, it does not take the random variance between subjects and individual differences in effect sizes into account (Frömer et al., 2018; Pernet et al., 2011). In order to account for this, we used the linear mixed effect models. The fixed effects included - additionally to the variables of interests (i.e., category and age group) - the baseline, interaction of learning category, age group and baseline, session, eye movements, gender and the random effects included subject, stimulus number, repetition number and sequence number.

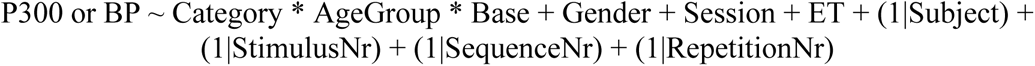

However, the linear mixed models do not provide test statistics for all learning states and group comparisons. Therefore, we computed the contrasts between the learning states and age groups using the *emmeans* package in R Studio and corrected the p-values for multiple comparisons using Bonferroni correction by multiplying p-values by the number of comparisons (Lenth, 2020). The adjusted p-values smaller than 0.05 (p < 0.05) were considered significant. The results of contrast comparisons were averaged over remaining fixed effects.

##### Finer temporal analysis - distinct functions of P300 and BP

To investigate distinct functions in memory formation theorized for P300 and BP we examined the finer temporal development of the P300 and BP amplitude. We expected the P300 amplitude to monotonically decrease from the first contact with the stimuli, as the surprise faded and the BP amplitude to peak at the point of the first accurate recall, where the stimulus was successfully committed to the memory and then gradually decrease. The stimuli were sorted based on the distance (number of sequence repetitions) to the first accurate recall of a given sequence element (i.e., newly learned point). Instead of having only 3 learning states (i.e., categories), we examined the development of P300 and BP amplitudes with respect to the point of the first accurate recall of a given sequence element (i.e., newly learned point) based on distance from this point. The stimuli categorized as newly learned have a distance of zero, whereas stimuli categorized as known have positive and unknown - negative distances. Due to insufficient number of stimuli, we considered only up to 2 repetitions prior (-2) and 4 sequence repetitions following (+4) the point of first accurate recall. Thus, we were able to analyze the gradual changes in P300 and BP amplitude over repetition relative to first accurate recall and test the assumed different progressions of both signals.

##### Neural generators of P300 and BP

Next, to investigate whether the P300 and BP are generated by the same or different neural sources, we compared the topographies of P300 and BP using the nonparametric topographic analysis of variance (TANOVA; (Murray et al., 2008)). If the scalp topographies are similar, it is unlikely that they are driven by different neural generators (Koenig et al., 2011). In line with Murray et al. (2008), we first computed the participant’s average topographies for P300 and BP based on a time window between onset and offset of a component obtained using the Liesefeld (2018) method as detailed above. We then normalized the data by dividing all participant’s topographical maps by its global field power. Next, we computed the global dissimilarity index (DISS), which represents the difference between two electric fields. DISS can span between 0 and 2, where 0 means that topographical maps are identical and 2 indicates inverse topographies. However, as DISS is a single measure of the distance between two topographical maps, a non-parametric randomization statistical test must be performed in order to obtain an empirical distribution of DISS values under the null hypothesis. To do so, we permuted the data by randomly shuffling within subject the position of electrodes on each iteration for P300 and BP (in each iteration, the position of electrodes was changed in an identical way in all participants, but differently for P300 and BP), recomputing the group average topographical maps and recomputing the DISS. We repeated this procedure 1000 times. Finally, in order to obtain the p-value for this permutation test, we counted the number of DISS as or more extreme than the initial DISS and divided by the number of permutations (i.e., 1000).

Obtaining a significant p-value in TANOVA allows us to reject the null hypothesis that the differences in scalp topographies are due to noise alone. However, TANOVA is not able to prove equivalency. Therefore, to test the equivalence of both topographic maps, we additionally computed two one-sided equivalence tests (TOST; (Lakens, 2017)). The equivalence test was performed in RStudio using the TOSTpaired function from the TOSTER package (Lakens, 2017). Again, we computed the average scalp topography for each participant and normalized the data by dividing all participant’s topographical maps by its global field power. Subsequently, we performed the TOST on each of the 105 electrodes. The equivalence bound was set to the middle effect size of d = 0.5 (for the TOST procedure with bounds set to the smallest effect size of interest (SESOI) based on power analysis see Supplementary Figure 4). The TOST provided t-value and p-value for lower and upper bound, and each of the larger of the two p-values and corresponding t-values were plotted on a topoplot to visualize the results. Significant p-values allow us to reject the null hypothesis meaning that the potentials on a given electrode are equivalent.

Finally, we investigated the neural generators of P300 and BP using source level analysis. The forward model was derived from the MNI ICBM 2009 template brain (https://www.bic.mni.mcgill.ca/ServicesAtlases/ICBM152NLin2009) using the OpenMEEG implementation (Gramfort et al., 2010) of the three-layer Boundary Element Method (BEM). A surface based cortical sheet source model with 2002 vertices was used. The spatial filters, which allow transforming the scalp sensor data to source estimates (i.e., inverse problem solution), were constructed by applying the minimum-norm estimation algorithm (Hämäläinen & Ilmoniemi, 1994). The noise covariance matrix required for this inverse operator was estimated from -100 to 0 ms prior to stimulus onset. Subsequently, the spatial filters were multiplied with time series data resulting in source estimates time series for each voxel. Next, individual P300 and BP source estimates were computed for each participant based on the individual time window between onset and offset of a given component, from which the P300 and BP grand average source estimates topographies were derived and visualized.

#### 2. 6. 3. Reliability of behavioral performance and P300 amplitude

High test-retest reliability of outcome measures is essential for future longitudinal studies and the precise identification of biomarkers. Therefore, in order to assess the reliability of the performance measures (i.e., time constant tau) and P300 amplitude within- and between-sessions, we computed the intraclass correlation coefficients (ICC) for tau and P300. For within-session reliability, we computed the degree of consistency (i.e., norm-referenced reliability; ICC (C, 1) in (McGraw & Wong, 1996)) for tau and P300 across the three sequences in each of two sessions. The P300 for each sequence was defined as the average amplitude over all stimuli in the first three sequence repetitions (i.e., 24 stimuli). Sensitivity analysis revealed that the latter approach yielded equivalent results, when defining P300 as a sum over all stimuli in a given repetition. However, this would result in a different number of stimuli per participant, therefore we opted for the first approach for defining P300 (i.e., the average amplitude over all stimuli in the first three sequence repetitions). For between-session reliability (i.e., test-retest reliability), we computed the degree of consistency for measurements that are averages of three measurements (i.e. three sequences per session) of randomly selected participants (ICC (C, k) in (McGraw & Wong, 1996)). First, we computed the average tau and average P300 for each participant in a given session. Tau was defined as the average tau of three sequences in a given session and the P300 was defined as the average amplitude over all stimuli in the first three sequence repetitions and over all three sequences in a given session. Based on (Cicchetti, 1994), ICC less than 0.40 were considered poor, ICC between 0.40 and 0.59 fair, ICC between 0.60 and 0.74 good and ICC between 0.75 and 1.00 excellent.

## 3. Results

### 3. 1. Behavioral data

#### 3. 1. 1. Demographics

Table 1 shows basic demographic information for young and older participants.

#### 3. 1. 2. Visual sequence learning task performance

Based on the participants’ manual responses in the test phase of the sequence learning task, each stimulus was classified to one of the following categories: unknown, newly learned and known, resulting in an average per subject of 25.89 ± 5.56 unknown, 49.61 ± 1.59 newly learned and 120.97 ± 4.72 known stimuli in young and 67.15 ± 7.68 unknown, 60.16 ± 2.99 newly learned and 137.57 ± 6.91 known stimuli in older individuals (M ± SD). The development of the number of trials in each stimulus category over sequence repetition is depicted on Figure 1C.

We next tested the hypothesis that the knowledge about sequence elements increases with each sequence repetition and tested for potential age differences. Figure 2 shows the progression of knowledge index and learning index (see section 2.4) over the sequence repetition. The apparent decrease of knowledge index after the fourth sequence repetition in young participants can be explained by the fact that some subjects completed the task after four sequence repetition and the average knowledge index from this sequence repetition on represents the average over the remaining, poorer-performing participants. Therefore, we further subdivided the participants into groups based on the number of sequence repetitions required for memorization of all sequence elements. The black lines represent the learning curves of each of those groups. This subdivision indicates that in fact there is no decrease of knowledge over the repetitions.

**Figure 2.**
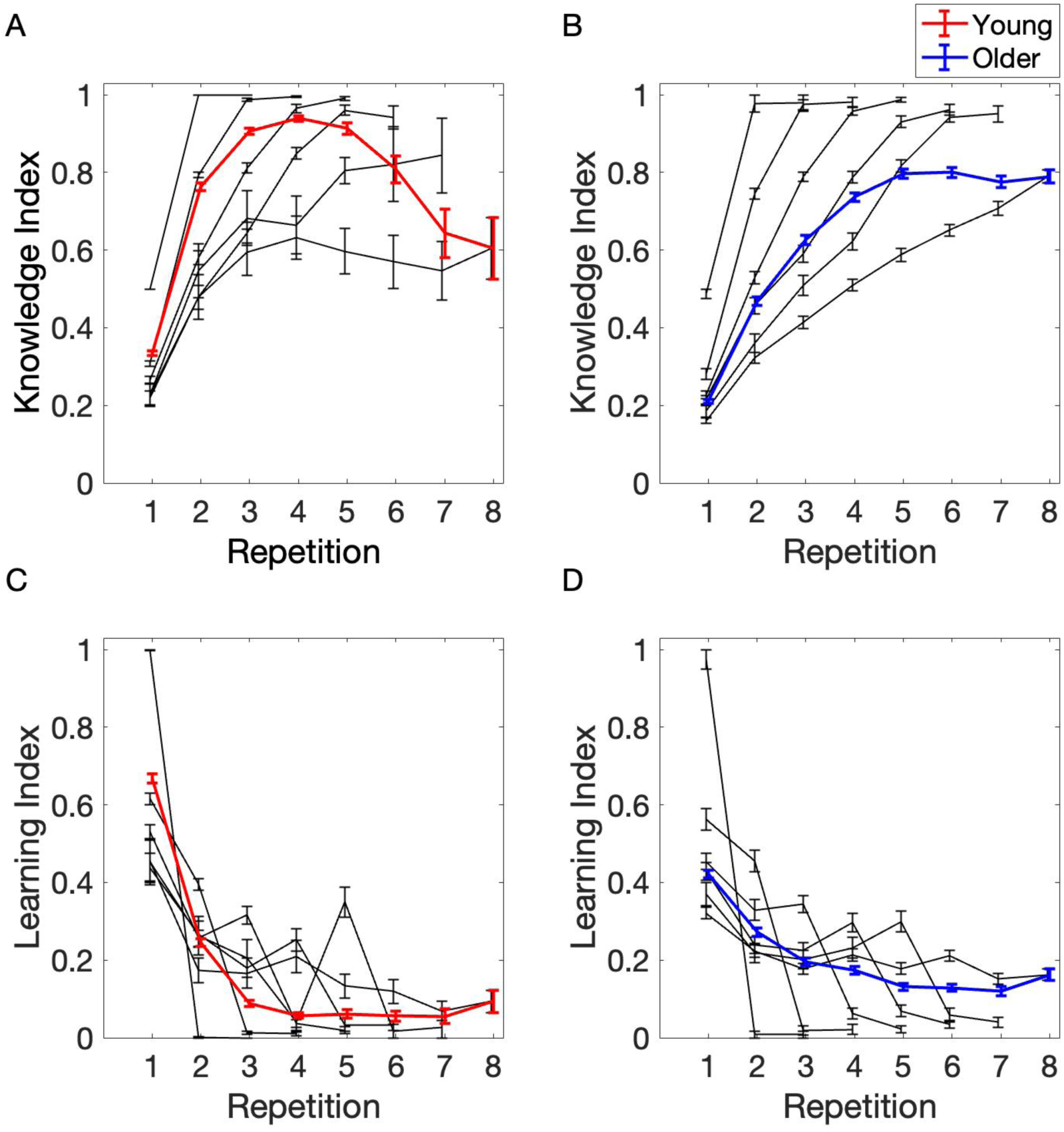
Average knowledge index (A-B) and learning index (C-D) across sequence repetitions in young (red) and older (blue) participants. The task lasted 3 to 8 repetitions, depending on learning speed. The black lines indicate the learning curves of participants that completed the task after 3, 4, 5, 6, 7 or 8 sequence repetitions. After 4th repetition the average knowledge index seems to decrease as the average is only computed for the remaining participants (for a KI, P300 and BP in each of the six sequences see Supplementary Figure 2). Error bars represent the standard error of the mean.

##### Knowledge Index

The best-fit model for the knowledge index included the fixed effect of repetition number, age group, and their interaction as well as the random effect of subject and sequence number (Table 2).

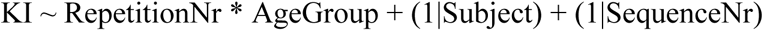

**Table 2.**
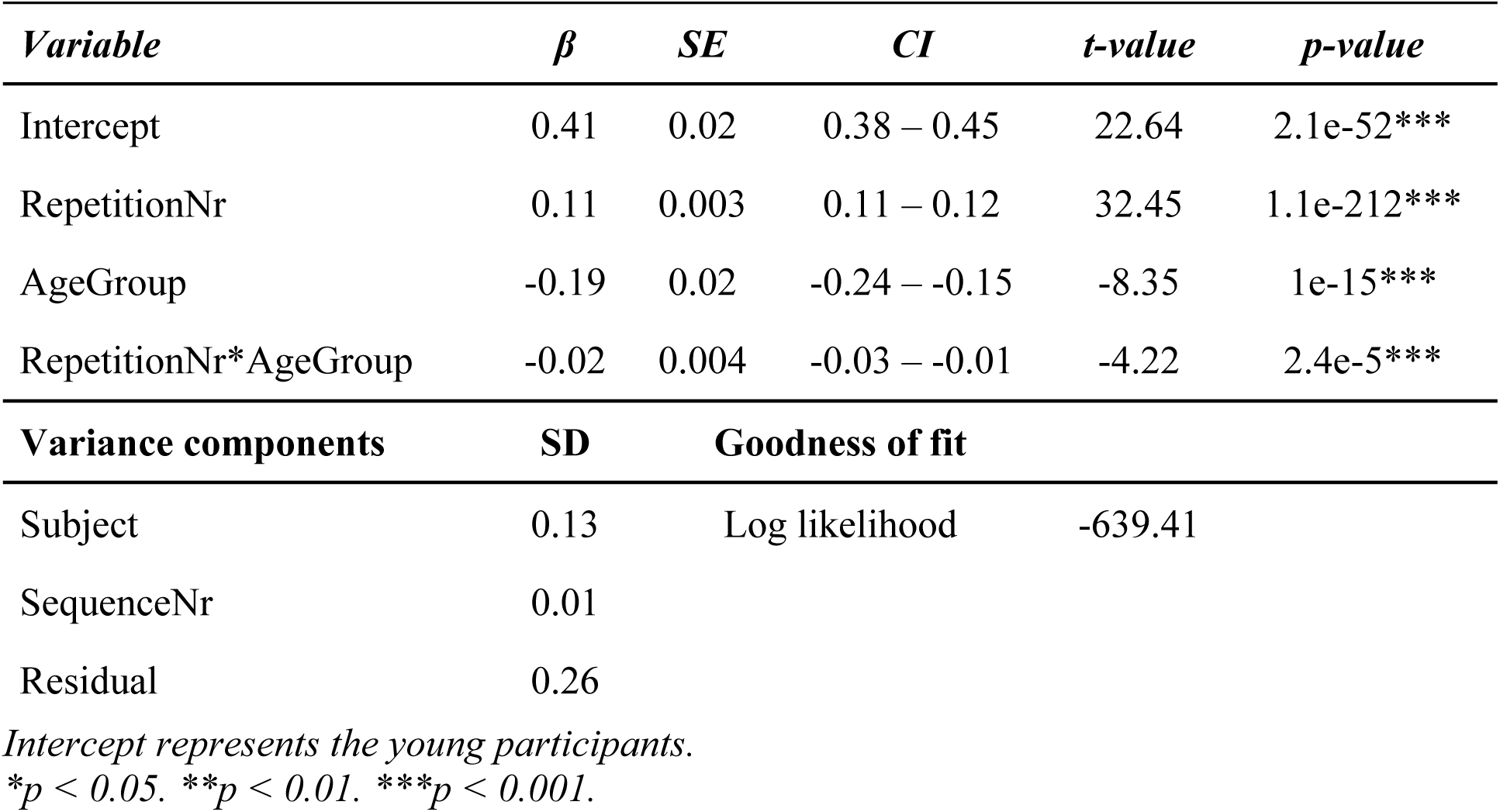
Effects of repetition number and age group on knowledge index

The model revealed a significant main effect of repetition number (β = 0.11, CI = [0.11; 0.12], p = 1.1e-212) that is the *knowledge index (KI)* increased with each sequence repetition (Figure 2A-B). Moreover, there was a significant main effect of age group (β = -0.19, CI = [-0.24; - 0.15], p = 1e-15), suggesting a lower *knowledge index* in the older group. There was also a significant interaction of repetition number and age group, indicating a smaller increase in *knowledge index* with each sequence repetition in the older group compared to the increase in the young group (β = -0.02, CI = [-0.03; -0.01], p = 2.4e-5). In addition, we observed a substantial variation of KI between subjects (SD = 0.13) and sequences (SD = 0.01).

##### Learning Index

The best-fit model for the *learning index (LI)* included only fixed effects: the repetition number, age group and their interaction (Table 3). The step function eliminated all random effects suggesting no substantial variance in the LI between participants or sequences. Therefore, subsequent results represent the coefficients of a simple linear model.

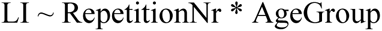

**Table 3.**
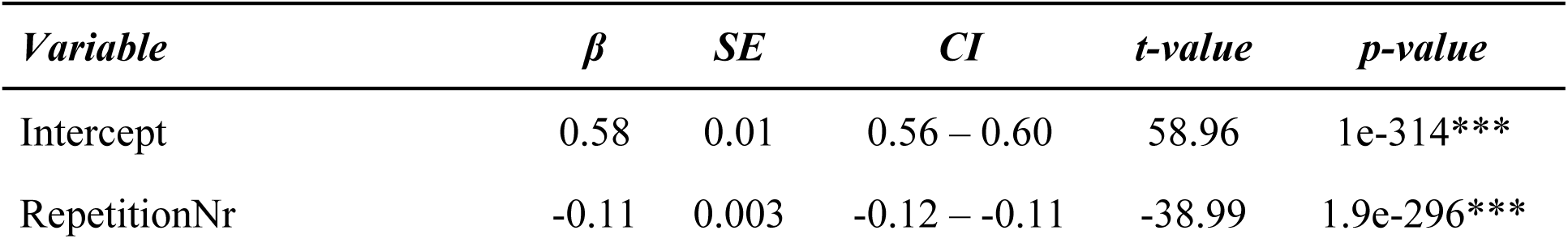
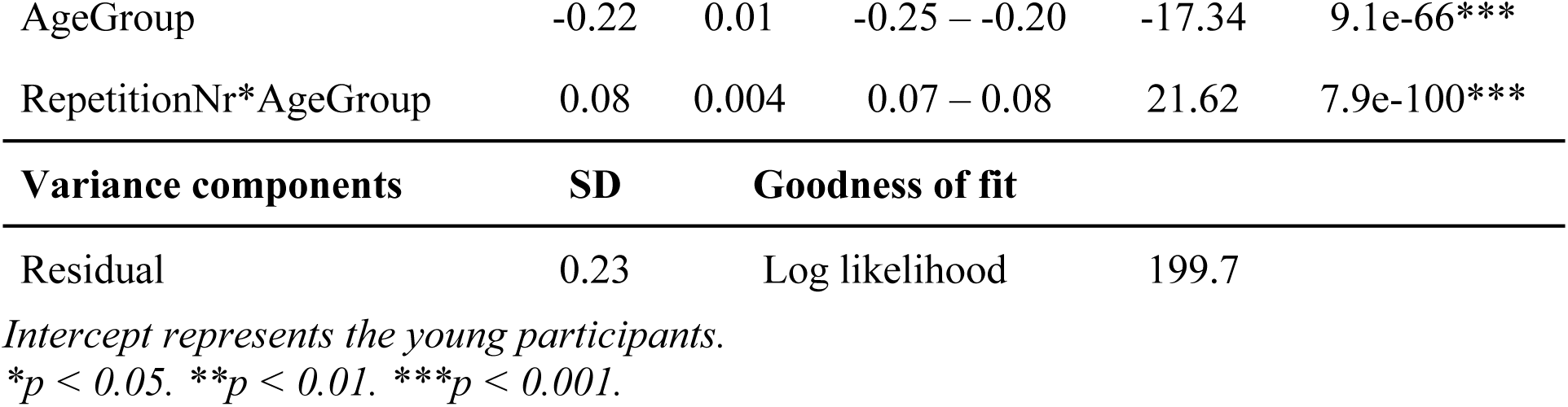
Effects of repetition number and age group on learning index

| *Variable* | $\beta$ | *SE* | *CI* | *t-value* | *p-value* |
| --- | --- | --- | --- | --- | --- |
| Intercept | 0.58 | 0.01 | 0.56 – 0.60 | 58.96 | 1e-314*** |
| RepetitionNr | -0.11 | 0.003 | -0.12 – -0.11 | -38.99 | 1.9e-296*** |

| AgeGroup | -0.22 | 0.01 | -0.25 – -0.20 | -17.34 | 9.1e-66*** |
| --- | --- | --- | --- | --- | --- |
| RepetitionNr*AgeGroup | 0.08 | 0.004 | 0.07 – 0.08 | 21.62 | 7.9e-100*** |
| Variance components | SD | Goodness of fit |  |  |  |
| Residual | 0.23 | Log likelihood | 199.7 |  |  |

*Intercept represents the young participants.*
\* $p < 0.05$ . \*\* $p < 0.01$ . \*\*\* $p < 0.001$ .

There was a significant main effect of repetition number (β = -0.11, CI = [-0.12; -0.11], p = 1.9e-296), meaning that the *learning index* decreased with each sequence repetition (Figure 2C- D). The learning index decreased, because with an increasing number of known stimuli, less stimuli that could be newly learned were available. In addition, there was a significant main effect of the age group, indicating a lower learning index in the older group (β = -0.22, CI = [- 0.25; -0.20], p = 9.1e-66) than in the young group. We further observed a significant interaction of repetition number and age group, reflecting the fact that the learning index decreased less strongly with each sequence repetition in the older group (β = 0.08, CI = [0.07; 0.08], p = 7.9e- 100). Thus, older participants tended to learn fewer sequence items in the first and second repetition, so that their learning was more evenly spread across repetitions.

### 3. 2. Neurophysiological data

#### 3. 2. 1. Interindividual variability of P300 peak latency

We began the analysis of neurophysiological data by exploring the interindividual variability of the P300 peak latency, because the potential amplitude differences between age groups might be overestimated using fixed time windows for extraction of P300. Analyzing the P300 and BP components using the same fixed time window for different age groups might reveal age-related alterations solely driven by latency differences. In addition, if older subjects exhibit an increased variability of P300 peak latencies with advancing age, this might introduce a bias to underestimate the true P300 amplitude in older individuals, even when using a fixed age-adjusted time window, but not accounting for inter-individual differences in P300 latency. Therefore, we test statistically the differences in P300 peak variability using the Levene’s test. The small p-value (Levene’s (1, 212) = 33.79, p = 2.2e-8) indicates that the variances of both age groups are not equal. Further inspection of the data revealed greater interindividual variability of P300 peaks in older participants. Figure 3B shows the distribution of individual P300 peaks in young (red) and older (blue) individuals. These results highlight the importance of computing P300 peaks individually, especially when investigating different age groups.

#### 3. 2. 2. P300 and BP amplitude across sequence repetition

So far, we established that young and older participants gradually learn the stimuli over repeated sequence presentations, with the young learning on average faster than older participants. In the next step, we tested the hypothesis that the learning progress is accompanied by decreasing P300 (due to increasing expectancy) and BP amplitudes (due to the decreasing need for active memory formation). For that, we computed the average P300 (mP300) and BP (mBP) amplitudes in each sequence repetition (i.e., average P300 and BP amplitude over eight stimuli in a given sequence repetition). Please note, some participants completed the task after just three sequence repetitions, while others needed up to eight. Consequently, the number of overall sequence repetitions decreases after the third repetition (see Figure 2). The average ERP waveforms in each sequence repetition are presented on Figure 4. Panel A shows the ERPs across sequence repetitions in young participants, and panel B shows the ERPs in older participants.

**Figure 4.**
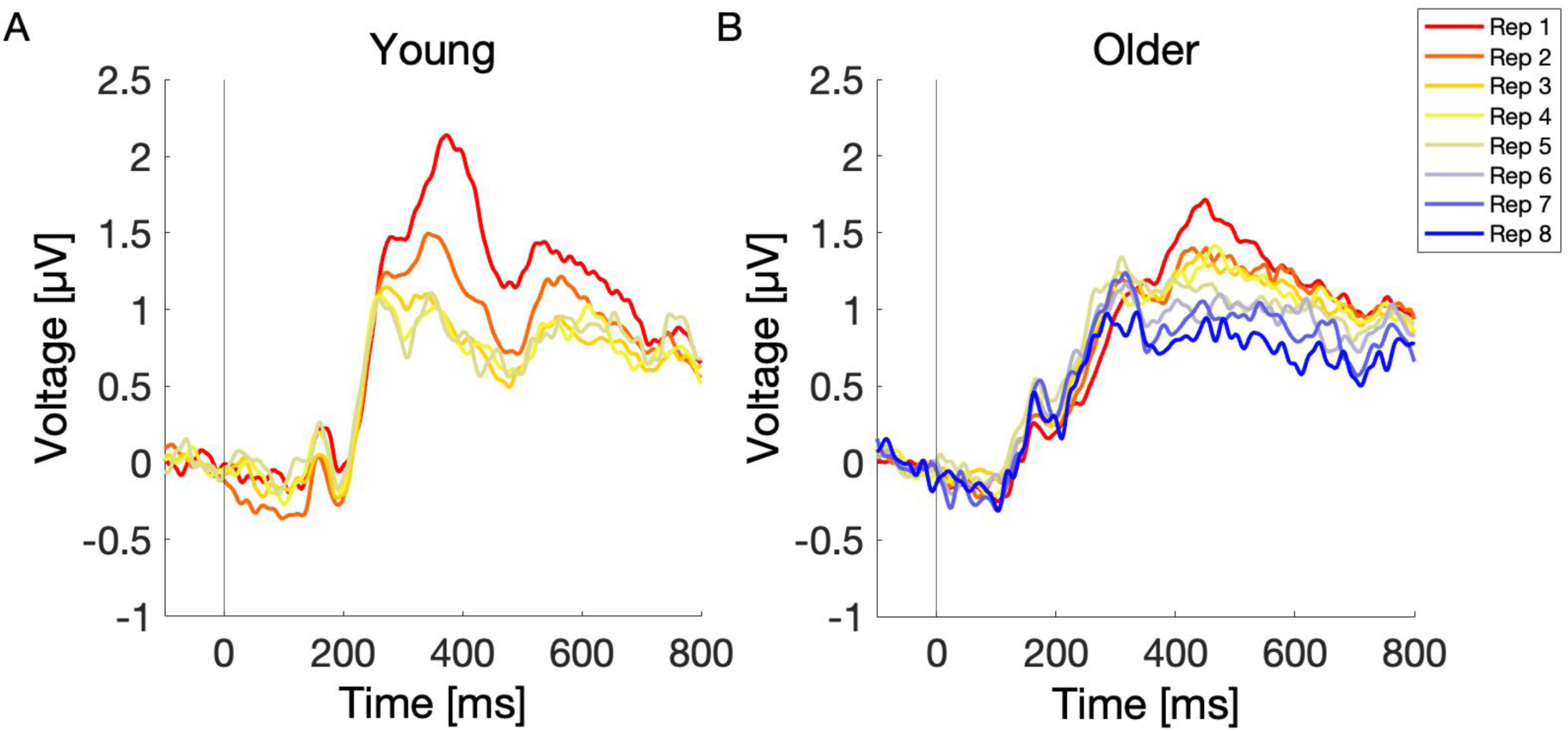
Average ERPs across sequence repetitions in young (A) and older (B) individuals. P300 and BP decreased significantly over sequence repetition in both, young and older participants. In young, only the first five sequence repetitions were plotted due to low amount of stimuli from the sixth sequence repetition on (only for plotting purposes). For a topographical maps of P300 and BP for each repetition see Supplementary Figure 5.

##### P300

The best-fit model for average P300 amplitude included the fixed effects of repetition number, age group, baseline, interaction of repetition number and age group, interaction of repetition number and baseline. The random part consisted of the subject and sequence number (Table 4).

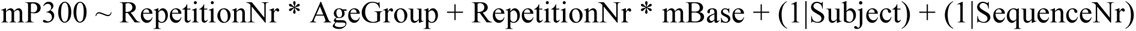

The model revealed a significant main effect of repetition number (β = -0.19, CI = [-0.21; - 0.16], p = 1.6e-39), indicating a decrease of P300 amplitude with each sequence repetition. Furthermore, there was a significant main effect of age group (β = -0.24, CI = [-0.40; -0.08], p = 0.003), that is, the P300 amplitude was decreased in older compared to young participants. There was also a significant interaction of repetition number and age group (β = 0.11, CI = [0.08; 0.14], p = 1.8e-11), indicating a smaller decrease of P300 amplitude across sequence repetitions in older participants. In addition, we observed a substantial variation of P300 amplitude between subjects (SD = 0.42) and sequences (SD = 0.06).

**Table 4.**
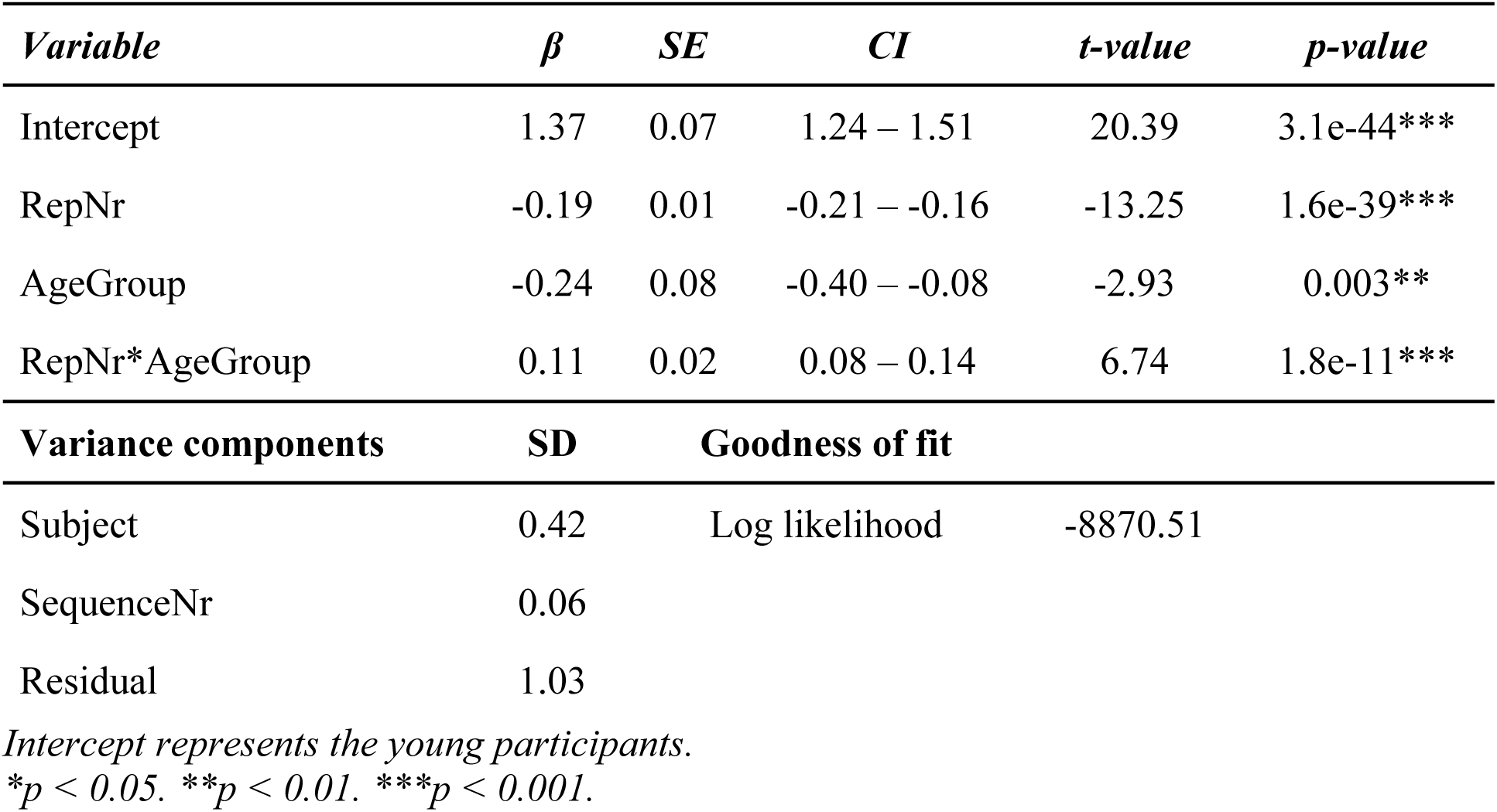
Effects of repetition number and age group on P300 amplitude

##### Broad Positivity

Subsequently, we identified the best-fit model for the average Broad Positivity amplitude across sequence repetitions (Table 5).

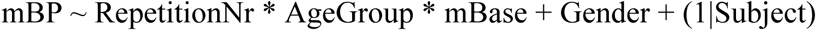

This model revealed a significant main effect of repetition number (β = -0.12, CI = [-0.15; - 0.09], p = 5.7e-18), indicating a decrease in BP amplitude with each sequence repetition. The main effect of age group (β = -0.03, CI = [-0.18; 0.13], p = 0.743) was not statistically significant, suggesting no age differences in overall BP amplitude aggregated across repetitions between young and older participants. However, there was a significant interaction of repetition number and age group (β = 0.04, CI = [0.01; 0.08], p = 0.007), reflecting a more gradual decrease of BP amplitude across sequence repetitions in older participants. Again, there was a considerable variation between subjects (SD = 0.34).

**Table 5.** Effects of repetition number and age group on BP amplitude

| *Variable* | *$\beta$* | *SE* | *CI* | *t-value* | *p-value* |
| --- | --- | --- | --- | --- | --- |
| Intercept | 0.95 | 0.07 | 0.82 – 1.08 | 14.31 | 1.1e-39*** |
| Repetition | -0.12 | 0.01 | -0.15 – -0.09 | -8.67 | 5.7e-18*** |
| AgeGroup | -0.03 | 0.08 | -0.18 – 0.13 | -0.33 | 0.743 |
| RepetitionNr*AgeGroup | 0.04 | 0.02 | 0.01 – 0.08 | 2.69 | 0.007** |
| Gender Female | 0.13 | 0.05 | 0.02 – 0.23 | 2.36 | 0.018* |
| **Variance components** | **SD** | **Goodness of fit** |  |  |  |
| Subject | 0.34 | Log likelihood |  | -8516.075 |  |
| Residual | 0.98 |  |  |  |  |

*Intercept represents the young participants.*
*\* $p < 0.05$ . \*\* $p < 0.01$ . \*\*\* $p < 0.001$ .*

#### 3. 2. 3. EEG signatures of successful learning

Having established that the P300 decreases with sequence repetition, we examined the potential of the P300 in predicting learning success across sequence repetitions, measured as knowledge index and learning index. For that, we computed the average P300 and baseline amplitude in each sequence repetition for each participant.

##### Knowledge Index and P300

Expectancy driven P300 was predicted to decrease, as the sequence knowledge strengthened. Thus, we tested whether the P300 amplitude could predict the knowledge index across sequence repetitions. First, we identified the best-fit model for predicting the knowledge index. The model included fixed effects of average P300 and age group. The random effects included only the subject (Table 6).

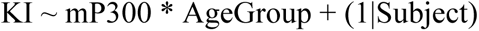

**Table 6.** Effects of P300 on Knowledge Index

| *Variable* | *$\beta$* | *SE* | *CI* | *t-value* | *p-value* |
| --- | --- | --- | --- | --- | --- |
| Intercept | 0.79 | 0.01 | 0.16 – 0.18 | 70.41 | 6e-210*** |
| mP300 | -0.07 | 0.006 | -0.08 – -0.06 | -12.92 | 1.1e-37*** |
| AgeGroup | -0.18 | 0.02 | -0.21 – -0.15 | -12.05 | 7.4e-28*** |
| mP300*AgeGroup | 0.03 | 0.007 | 0.02 – 0.05 | 4.71 | 2.5e-06*** |

| Variance components | SD | Goodness of fit |  |
| --- | --- | --- | --- |
| Subject | 0.08 | Log likelihood | -1684.67 |
| Residual | 0.31 |  |  |

*Intercept represents the young participants.*
*\* $p < 0.05$ . \*\* $p < 0.01$ . \*\*\* $p < 0.001$ .*

The model revealed a significant main effect of P300 (β = -0.07, CI = [-0.08; -0.06], p = 1.1e- 37) that is the knowledge index increased with decreasing P300 amplitude in young individuals. However, the sensitivity with which knowledge depended on P300 amplitude was lower in older participants, as indicated by the significant interaction of P300 and age group (β = 0.03, CI = [0.02; 0.05], p = 2.5e-06). Again, there was a considerable variation between subjects (SD = 0.08).

##### Learning Index and Broad Positivity

BP, the signal thought to reflect active memory trace formation, was expected to be maximal for stimuli being actively committed to memory. Thus, we tested whether the LI, which represents the proportion of newly learned stimuli in a given repetition, could be predicted by the average BP amplitude across sequence repetitions. First, we identified the best-fit model for predicting the learning index. The model included fixed effects of average BP, age group, average baseline, as well as their 3-way interaction (Table 7).

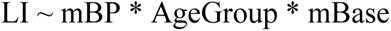

**Table 7.**
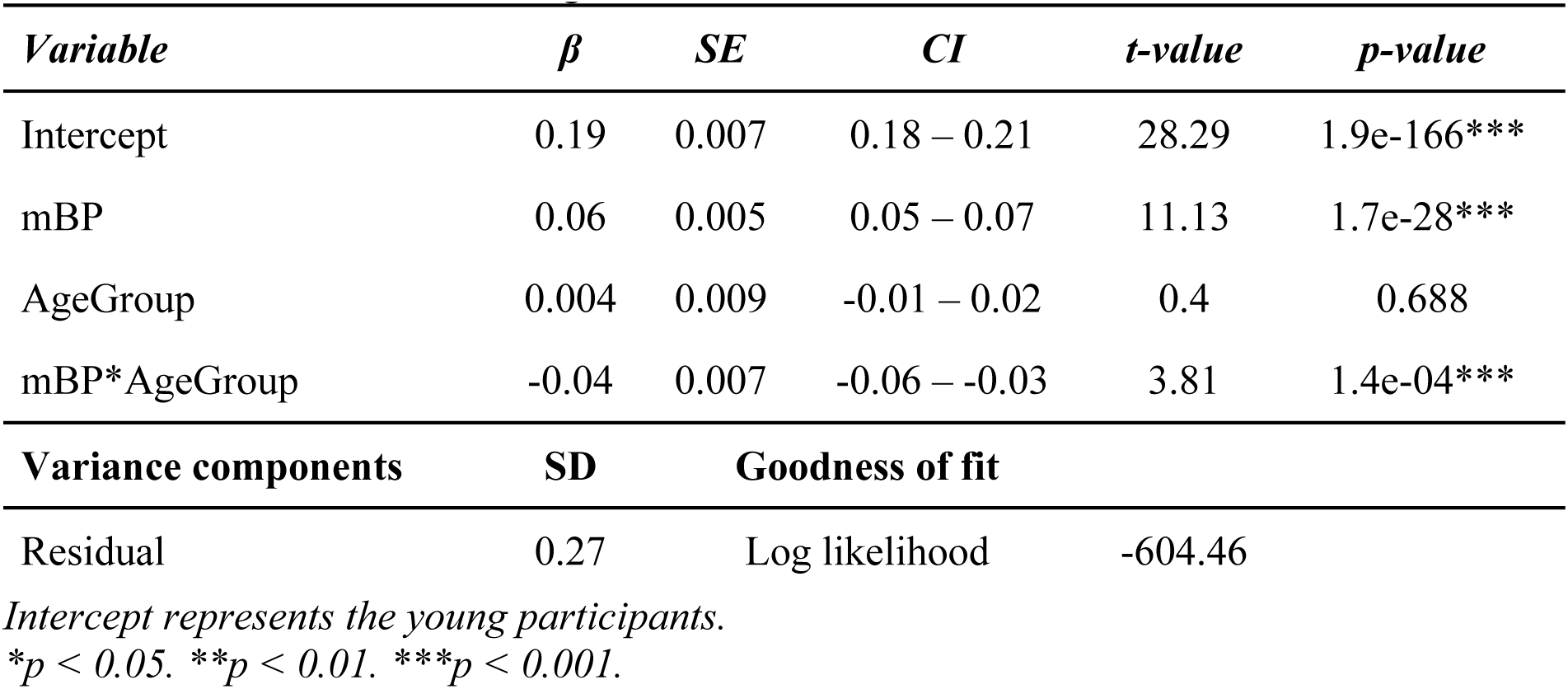
Effects of BP on Learning Index

The model revealed a significant main effect of the BP (β = 0.06, CI = [0.05; 0.07], p = 1.7e- 28), indicating a positive relationship between the learning index and BP. The main effect of the age group did not reach significance (β = 0.004, CI = [-0.01; 0.02], p = 0.688), but a significant interaction of age group and BP (β = -0.04, CI = [-0.06; -0.03], p = 1.4e-04) indicated that LI varied less sensitively with BP amplitude in the older group.

Previous analysis demonstrated that the P300 and BP were able to predict KI and LI, respectively, more strongly in young participants. The significant interaction of both components with age groups raised the question of whether the KI and LI could be predicted in older participants considered alone. Therefore, we computed two separate linear mixed models with KI and LI as dependent variables respectively, only for the older group. For KI, the model revealed a significant effect of average P300 (β = -0.04, CI = [-0.05; -0.03], p = 1.1e-12), indicating that KI can be predicted by the P300 amplitude within older participants. For LI, the model revealed a significant main effect of average BP (β = 0.01, CI = [0.01; 0.02], p = 2e-4), demonstrating that LI can also be predicted in older participants.

#### 3. 2. 4. Predicting learning success across participants

Finally, we aimed to predict the learning success across participants and to identify fast and slow learners based on the decrease of the P300 amplitude between the first and third sequence repetition. To facilitate the measure of learning success, we fitted an exponential function to each participant’s learning curve and obtained a time in sequence repetitions, here called tau, in which each participant learned 63% or 1 – 1/e of all sequence elements. Subsequently, we tried to predict the tau using the decrease of the P300 amplitude as a predictor. Figure 5A shows the average learning curve and average tau across all participants. Figure 5B shows the decrease of the P300 amplitude over sequence repetitions. The best-fit model included only the fixed main effects of P300 decrease and age group, interaction of age group and P300 decrease, as well as the random effect of the subject (Table 8). The model revealed a significant main effect of P300 decrease (β = 0.06, CI = [0.01; 0.10], p = 0.015) indicating greater tau (i.e., slower learning) with smaller P300 decrease. Moreover, the significant main effect of age group (β = 1.76, CI = [1.52; 2.16], p = 9e-25) showed greater tau in older compared to that of young participants, meaning that the older learned significantly slower than the young individuals. There was also a significant interaction of P300 decrease and age group (β = -0.07, CI = [-0.14; -0.01], p = 0.042), indicating that for the older participants the P300 decrease was less predictive of the time constant tau. Nevertheless, we further investigated the differences in slopes of young and older participants highlighted on Figure 5C-D. We computed a separate linear mixed effect model for young and older, with the P300 decrease as a fixed effect and the subject as a random effect. In young, the model revealed a significant effect of P300 decrease (β = 0.06, CI = [0.03; 0.09], p = 2.8e-4), confirming previous analysis. However, the P300 decrease failed to reach significance in older participants (β = -0.01, CI = [-0.08; 0.05], p = 0.663). Sensitivity analysis revealed that changing the definition of time constant tau (i.e., 30%, 50% or 70% instead of 63%) or computing the decrease of P300 amplitude from the first to second, or from the first to the fifth repetition, does not change the overall results.

**Figure 5.**
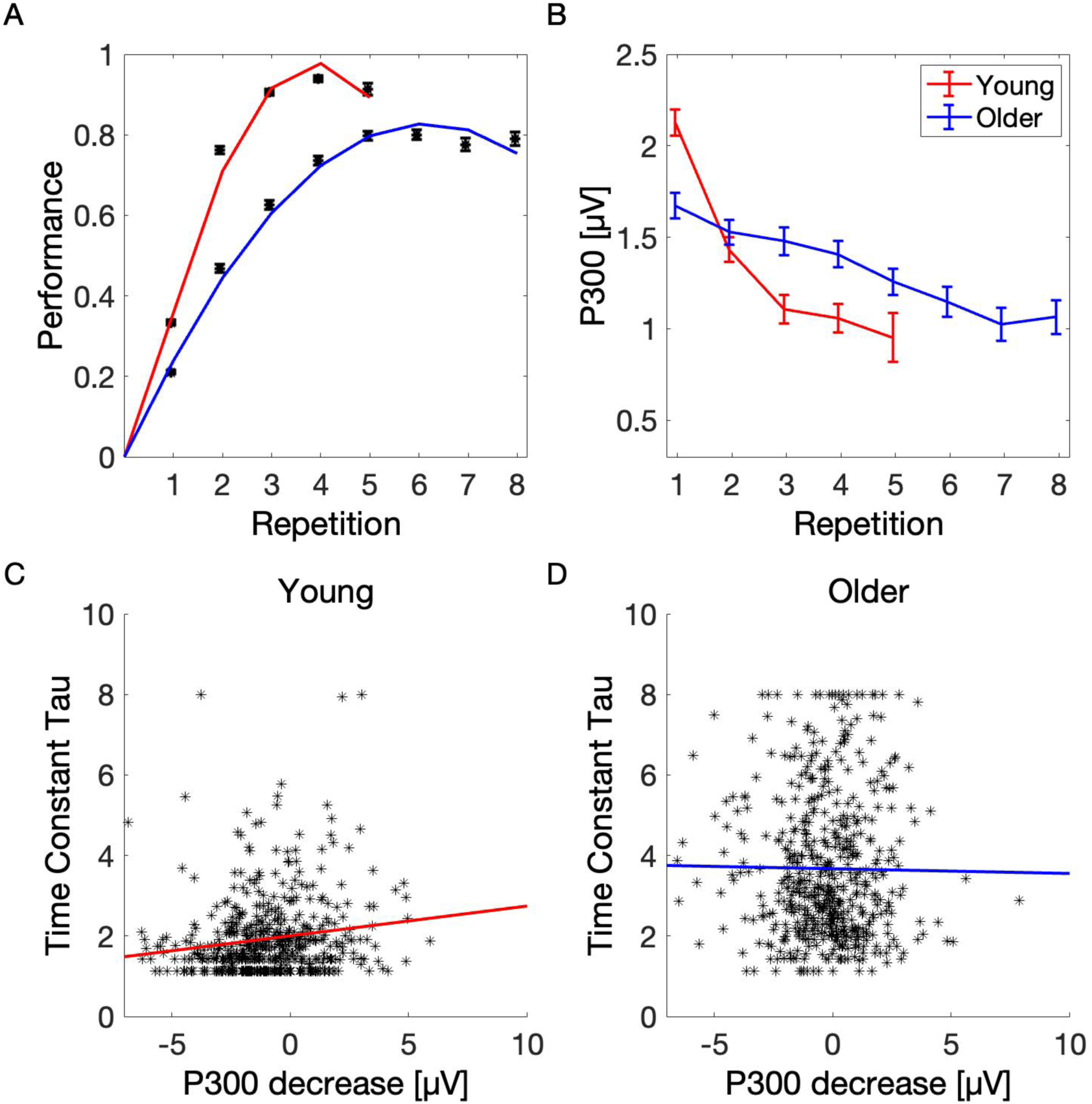
Learning performance and P300 decrease over sequence repetition. (A) A polynomial function was fitted to each participant’s learning curve in order to assess time (in repetitions), in which a participant has learned 63% of all sequence elements. Black error bars represent the average performance (i.e., knowledge index), and the red and blue curves represent the average polynomial function fitted to the performance in young and older participants. The dashed line depicts average tau in young (Tau = 1.8) and older (Tau = 3.02) participants. (B) Decrease of the P300 amplitude over sequence repetitions. In young participants, the repetitions 6 to 8 were not plotted due to insufficient number of stimuli. Error bars represent the standard error of the mean. (C-D) Prediction of learning success across young (C) and older (D) participants. We decided to compute the difference between the third and the first sequence repetition, where most of the learning took place. The decrease of P300 amplitude could significantly predict learning success (i.e., fast learners) in young, but not in older participants.

**Table 8.**
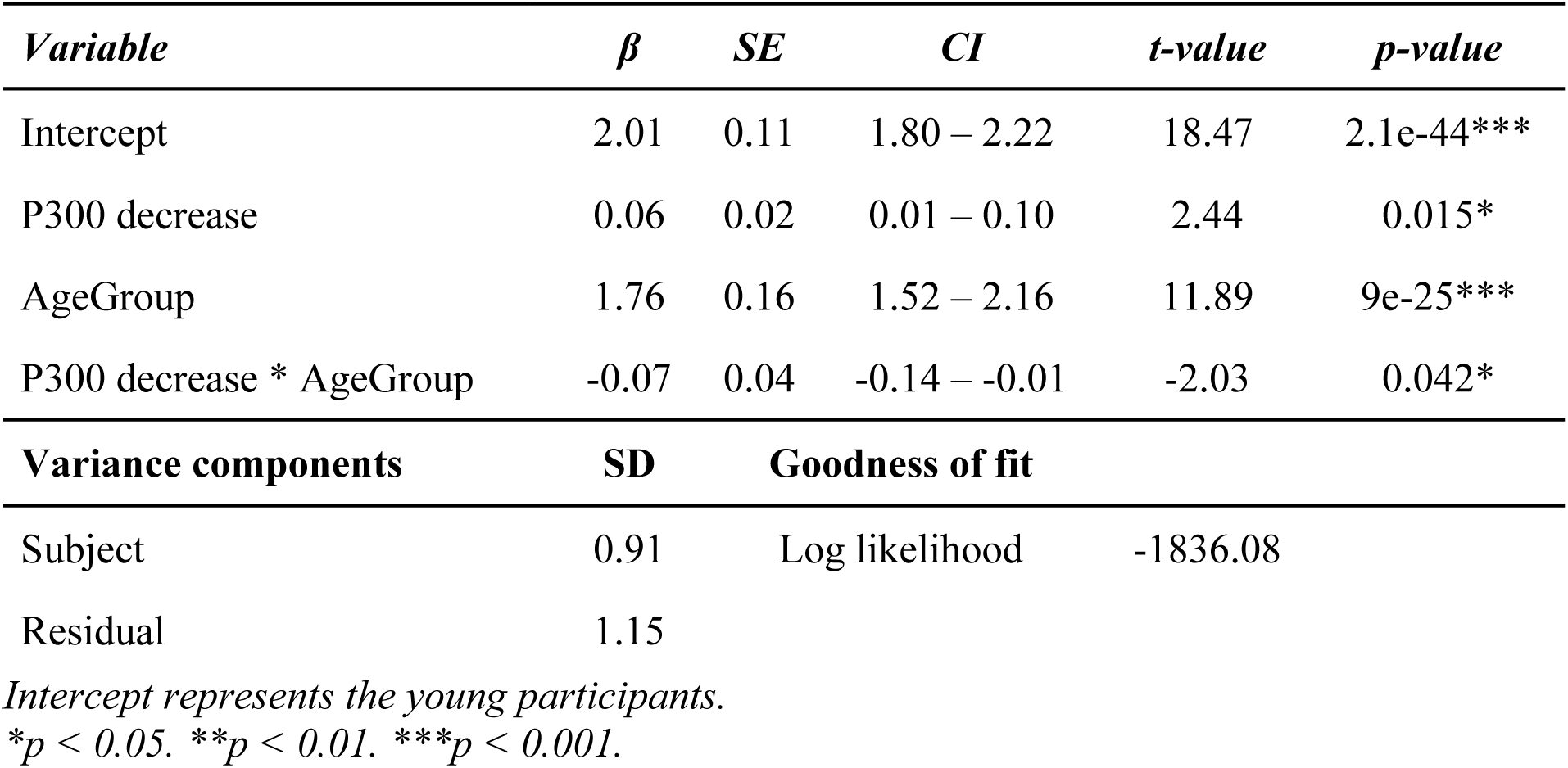
Effects of the P300 amplitude decrease on time constant tau.

#### 3. 2. 5. P300 and BP amplitude across learning categories and age groups

So far, we demonstrated that the P300 and BP amplitudes decrease with sequence repetitions and the decrease could predict learning success across repetitions and participants. However, the decrease of amplitude could be simply an effect of habituation and the component’s amplitude could decrease as a function of time spent on a task. Therefore, to provide stronger evidence for the relationship of P300 and BP with learning, we tested whether the amplitude of both components changes as a function of learning state, that is, from stimulus being unknown, to newly learned and finally fully known, regardless of the sequence repetition. First, we identified the best-fit model for the P300 and BP amplitude. Because we were merely interested in amplitude differences across learning categories and age groups, we additionally computed contrasts for those variables using the emmeans package in R Studio. Figure 6A-C shows the ERP waveforms and scalp topographies of young and older participants averaged over six centro-parietal electrodes as a function of the learning category. Figure 6D-E displays the P300 and BP amplitudes in young and older participants corrected for interindividual variability.

**Figure 6.**
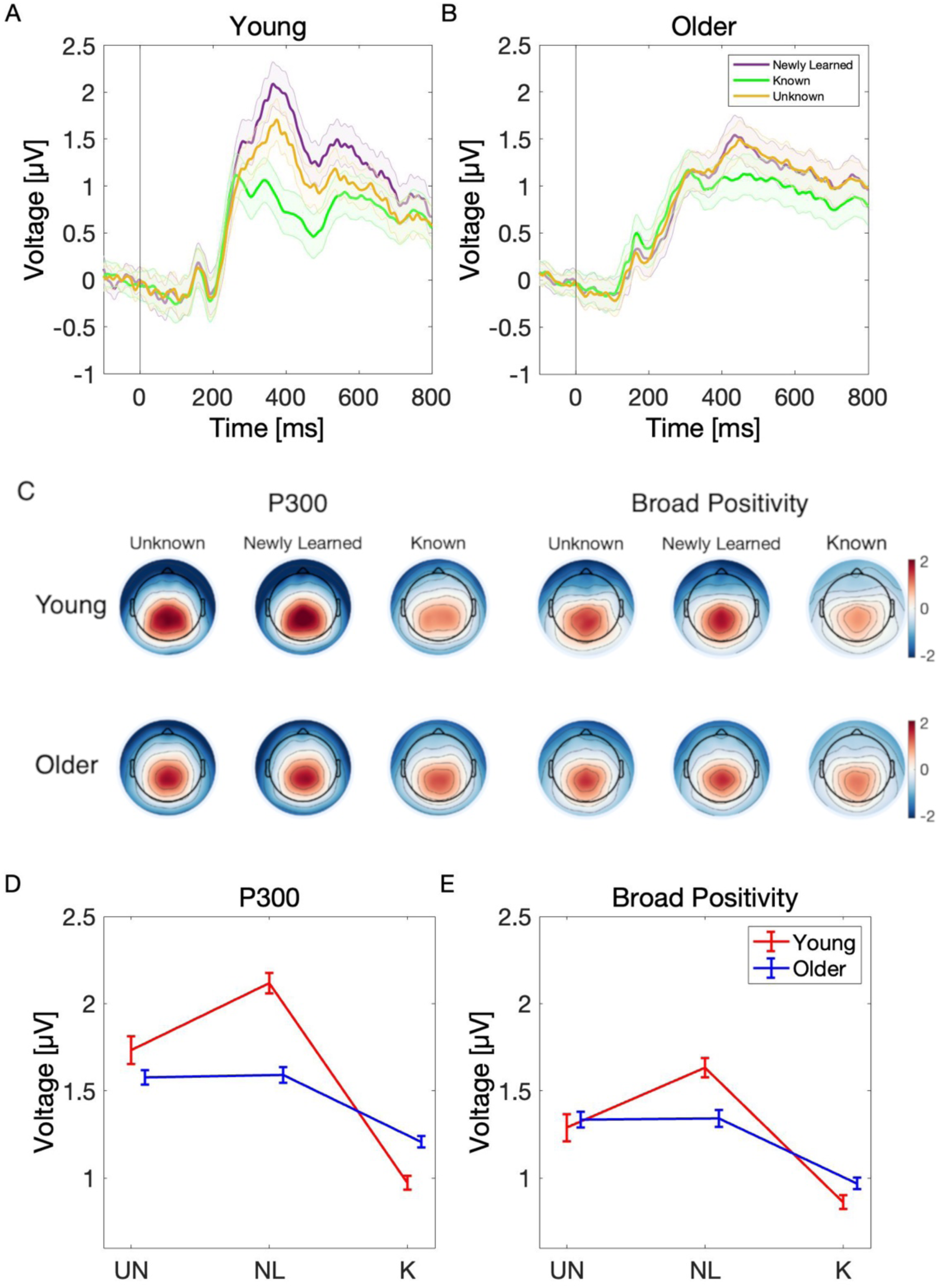
Event-related potentials, topographical maps and mean amplitudes over learning categories computed based on individual peaks. (A-B) Centro-parietal ERPs of young (left) and older (right) participants across learning categories: unknown, newly learned and known. Shaded error bars represent the standard error of the mean. (C) Scalp topographies of the P300 and broad positivity for each learning state and age group. (D-E) The linear mixed-effects model showed a significant interaction effect of category K and age group. The results of contrast comparisons revealed that the P300 amplitudes were greater for unknown and newly learned trials than for known trials in both age groups, however the amplitude difference between unknown and newly learned trials did not reach significance. Moreover, the P300 amplitude of newly learned trials was greater in young compared to older participants. The broader positivity showed the same effects during learning as the P300 within both age groups (UN = NL > K), however no age differences between contrasts were found. The errorbar represents the standard error of the mean.

##### P300

The best-fit model for P300 amplitude included the fixed effect of category, age group, baseline and their 3-way interaction, and random effects of subject, stimulus number, repetition number and sequence number (Table 9).

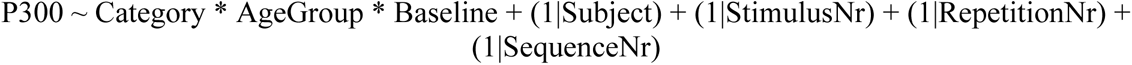

**Table 9.**
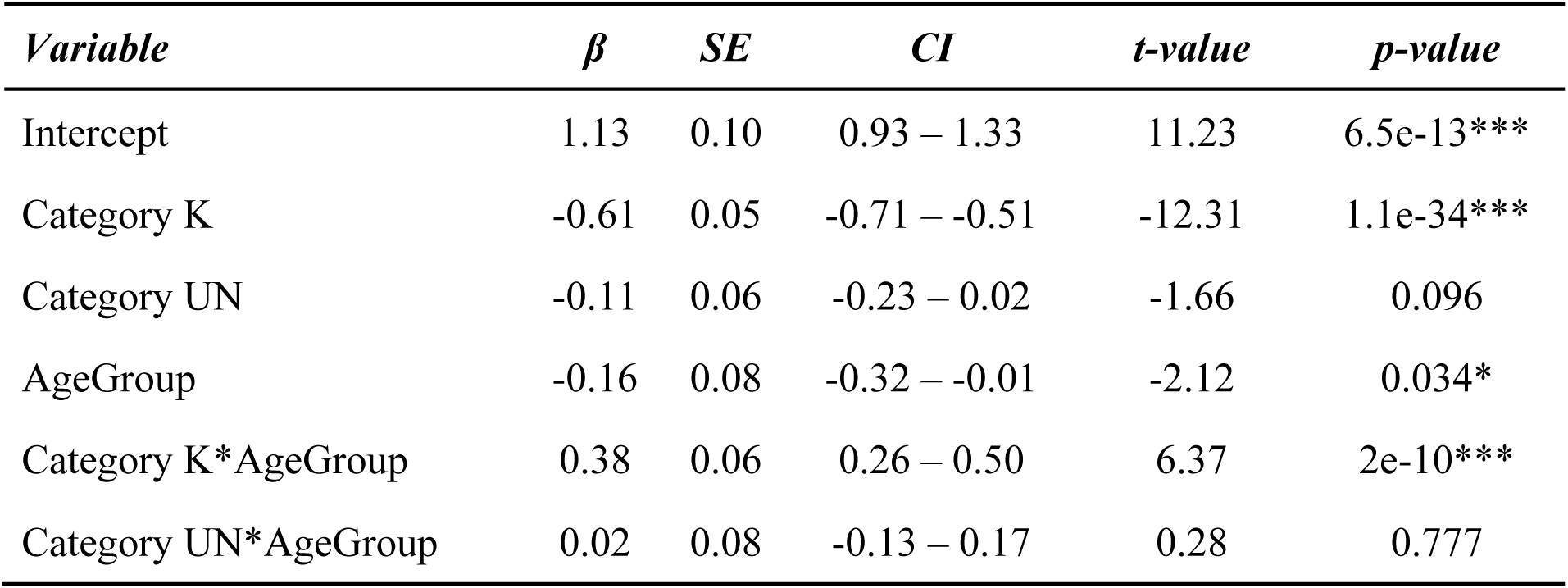
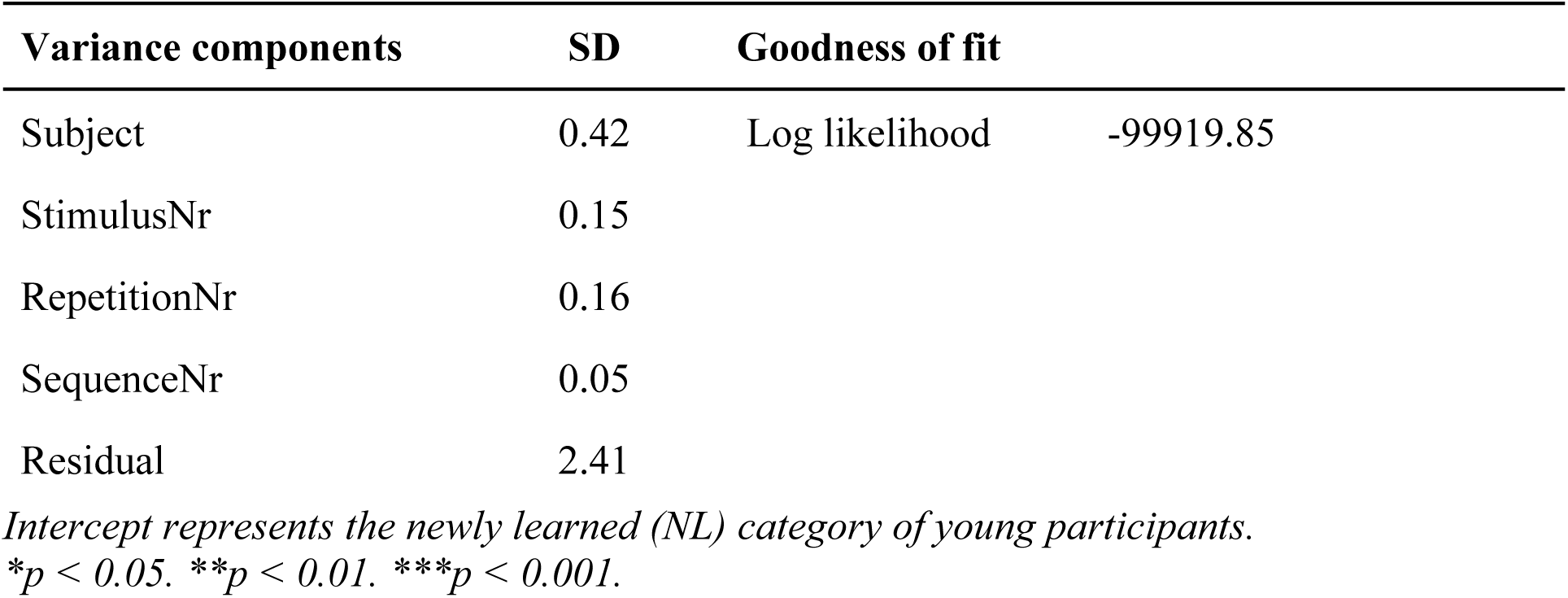
Effects of age group and learning categories on P300 amplitude

The model revealed a significant main effect of category K (β = -0.61, CI = [-0.71; -0.51], p = 1.1e-34), indicating decreased P300 amplitude for K compared to NL trials but a significant interaction of category K and age group (β = 0.38, CI = [0.26; 0.50], p = 2e-10) indicated that P300 amplitude of category K was less decreased in the older group. No other effects and interaction effects reached significance. Next, we moved on to computing contrasts between categories and age groups.

In young participants, the results of contrast comparisons showed a significantly greater P300 amplitude for UN compared to K (β = 0.49, CI = [0.37; 0.61], p = 1.7e-14) trials. Moreover, the P300 amplitude was significantly larger for NL than K (β = 0.65, CI = [0.55; 0.74], p = 3.1e-39) trials. The difference between NL and UN trials was not statistically significant (β = 0.16, CI = [0.04; 0.28], p = 0.09). Similarly, in older participants, the P300 amplitude was significantly greater in UN than K (β = 0.13, CI = [0.04; 0.21], p = 0.034) and NL than K (β = 0.20, CI = [0.13; 0.29], p = 5.8e-06) trials. The amplitude differences between NL and UN (β = 0.08, CI = [-0.01; 0.17], p = 0.623) did not reach significance. Finally, we compared the P300 amplitude in each learning category between age groups: there were no significant differences between age groups for the UN trials (β = 0.16, CI = [-0.01; 0.32], p = 0.509), but young participants had significantly greater P300 amplitude for the NL trials (β = 0.24, CI = [0.09; 0.30], p = 0.017) and significantly smaller amplitude for the K trials (β = -0.20, CI = [-0.33; - 0.07], p = 0.024). The results of contrast comparisons are summarized in Table 10 and visualized in Figure 6D.

**Table 10.**
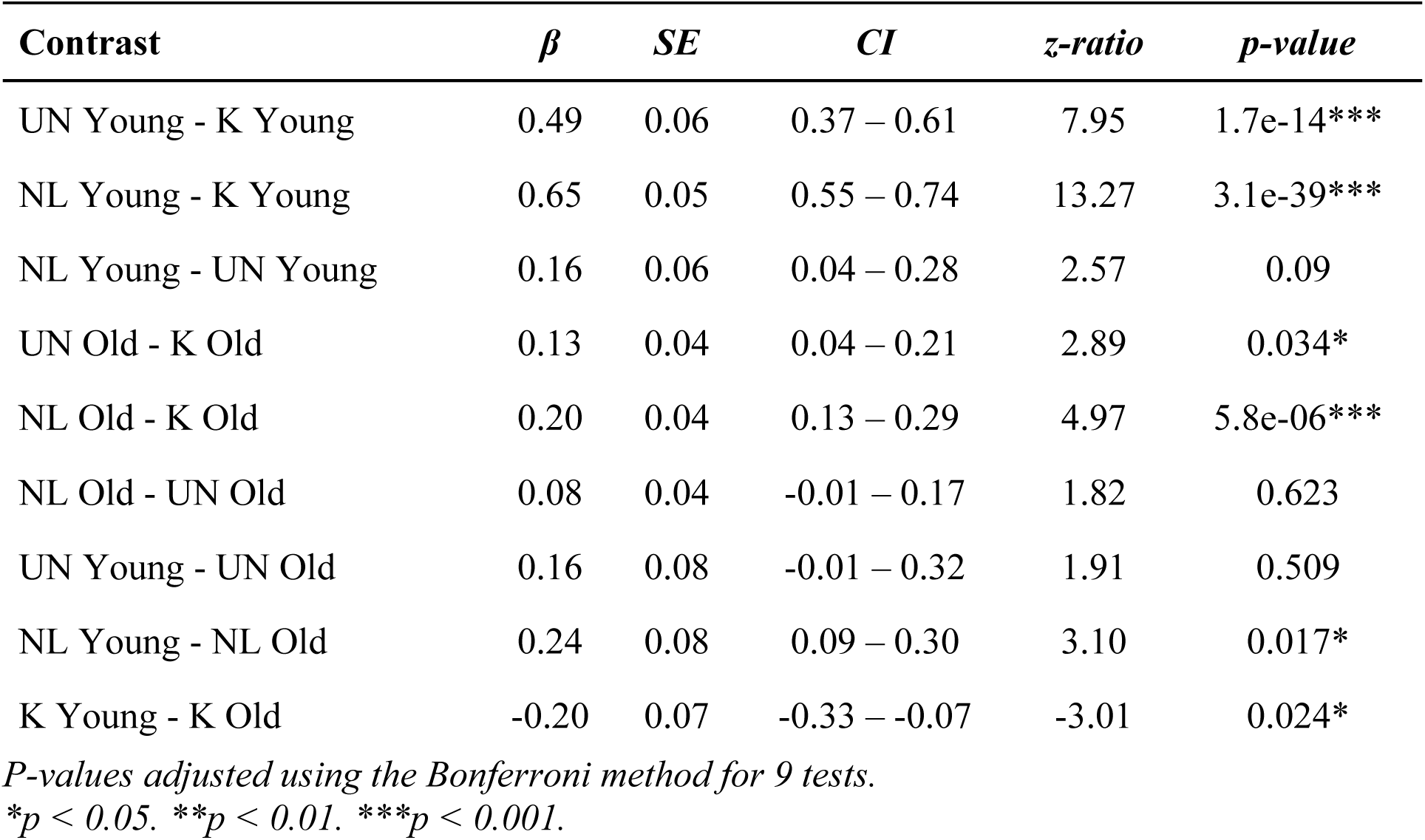
P300: contrast comparisons between age groups and learning categories.

| Contrast | $\beta$ | SE | CI | z-ratio | p-value |
| --- | --- | --- | --- | --- | --- |
| UN Young - K Young | 0.49 | 0.06 | 0.37 – 0.61 | 7.95 | 1.7e-14*** |
| NL Young - K Young | 0.65 | 0.05 | 0.55 – 0.74 | 13.27 | 3.1e-39*** |
| NL Young - UN Young | 0.16 | 0.06 | 0.04 – 0.28 | 2.57 | 0.09 |
| UN Old - K Old | 0.13 | 0.04 | 0.04 – 0.21 | 2.89 | 0.034* |
| NL Old - K Old | 0.20 | 0.04 | 0.13 – 0.29 | 4.97 | 5.8e-06*** |
| NL Old - UN Old | 0.08 | 0.04 | -0.01 – 0.17 | 1.82 | 0.623 |
| UN Young - UN Old | 0.16 | 0.08 | -0.01 – 0.32 | 1.91 | 0.509 |
| NL Young - NL Old | 0.24 | 0.08 | 0.09 – 0.30 | 3.10 | 0.017* |
| K Young - K Old | -0.20 | 0.07 | -0.33 – -0.07 | -3.01 | 0.024* |

*P-values adjusted using the Bonferroni method for 9 tests.*
*\* $p < 0.05$ . \*\* $p < 0.01$ . \*\*\* $p < 0.001$ .*

##### Broad Positivity

Next, we investigated the broader centro-parietal positivity. The best-fit model included fixed effects of category, age group, baseline, gender and an interaction of category, age group and baseline and the random effect of subject, stimulus number and repetition number (Table 11).

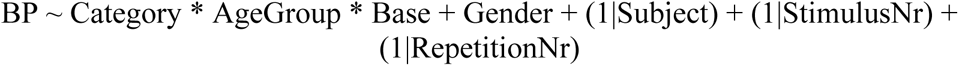

**Table 11.** Effects of age group and learning categories on BP amplitude

| *Variable* | $\beta$ | *SE* | *CI* | *t-value* | *p-value* |
| --- | --- | --- | --- | --- | --- |
| Intercept | 0.85 | 0.08 | 0.70 – 1.00 | 11.19 | 2.6e-17*** |
| Category K | -0.45 | 0.05 | -0.54 – -0.35 | -9.30 | 2.6e-20*** |
| Category UN | -0.13 | 0.06 | -0.25 – -0.01 | -2.04 | 0.041* |
| AgeGroup | -0.09 | 0.07 | -0.22 – 0.04 | -1.34 | 0.181 |
| Category K*AgeGroup | 0.17 | 0.06 | 0.06 – 0.29 | 2.97 | 0.003** |
| Category UN*AgeGroup | 0.10 | 0.08 | -0.05 – 0.25 | 1.34 | 0.181 |
| Gender Female | 0.11 | 0.05 | 0.02 – 0.21 | 2.26 | 0.024* |
| **Variance components** | **SD** | **Goodness of fit** |  |  |  |
| Subject | 0.33 | Log likelihood |  | -99090.4 |  |
| StimulusNr | 0.10 |  |  |  |  |
| RepetitionNr | 0.08 |  |  |  |  |
| Residual | 2.37 |  |  |  |  |

*Intercept represents the newly learned (NL) category of young participants.*
*\* $p < 0.05$ . \*\* $p < 0.01$ . \*\*\* $p < 0.001$ .*

The model revealed a significant main effect of category K (β = -0.45, CI = [-0.54; -0.35], p = 2.6e-20), indicating decreased BP amplitude for K compared to NL trials but a significant interaction of category K and age group (β = 0.17, CI = [0.06; 0.29], p = 0.003) indicated that BP amplitude of category K was less decreased in the older group. Further, we observed a significant main effect of category UN (β = -0.13, CI = [-0.25; -0.01], p = 0.041), suggesting decreased BP amplitude for UN compared to NL trials and a significant main effect of gender (β = 0.11, CI = [0.02; 0.21], p = 0.024), indicating greater BP amplitude in females compared to males. No other effects and interaction effects reached significance. Next, we moved on to computing contrasts between categories and age groups.

In young participants, the contrast comparison revealed a significantly larger BP amplitude for UN relative to K (β = 0.30, CI = [0.18; 0.41], p = 6e-6) and NL relative to K (β = 0.45, CI = [0.35; 0.54], p = 3.9e-20) trials. The BP amplitude of NL and UN trails was not statistically different (β = 0.15, CI = [0.03; 0.27], p = 0.127). Similarly, in older participants, we observed a larger amplitude for UN than K (β = 0.26, CI = [0.17; 0.34], p = 6.2e-09) and NL than K (β = 0.27, CI = [0.19; 0.35], p = 1.5e-10) trials, while the difference between UN and NL trials did not reach significance (β = 0.01, CI = [-0.07; 0.10], p = 0.999). When comparing both age groups, we found no differences in the UN (β = -0.03, CI = [-0.17; 0.11], p = 0.999), NL (β = 0.10, CI = [-0.03; 0.23], p = 0.999) and K (β = -0.07, CI = [-0.18; 0.04], p = 0.999) trials. The results of contrast comparisons are summarized in Table 12 and visualized in Figure 6E.

**Table 12.** Broad Positivity: contrast comparisons between age groups and learning categories.

| **Contrast** | $\beta$ | *SE* | *CI* | *z-ratio* | *p-value* |
| --- | --- | --- | --- | --- | --- |
| UN Young - K Young | 0.30 | 0.06 | 0.18 – 0.41 | 4.97 | 6e-6*** |
| NL Young - K Young | 0.45 | 0.05 | 0.35 – 0.54 | 9.43 | 3.9e-20*** |
| NL Young - UN Young | 0.15 | 0.06 | 0.03 – 0.27 | 2.45 | 0.127 |
| UN Old - K Old | 0.26 | 0.04 | 0.17 – 0.34 | 6.17 | 6.2e-09*** |
| NL Old - K Old | 0.27 | 0.04 | 0.19 – 0.35 | 6.74 | 1.5e-10*** |
| NL Old - UN Old | 0.01 | 0.04 | -0.07 – 0.10 | 0.31 | 0.999 |
| UN Young - UN Old | -0.03 | 0.07 | -0.17 – 0.11 | -0.44 | 0.999 |
| NL Young - NL Old | 0.10 | 0.07 | -0.03 – 0.23 | 1.66 | 0.999 |
| K Young - K Old | -0.07 | 0.06 | -0.18 – 0.04 | -1.23 | 0.999 |

*Results are averaged over the levels of Gender. P-values adjusted using the Bonferroni method for 9 tests.*
*\* $p < 0.05$ . \*\* $p < 0.01$ . \*\*\* $p < 0.001$ .*

#### 3. 2. 6. Finer temporal analysis - distinct functions of P300 and BP

So far, we established that the P300 and BP amplitudes change over the course of learning, from a sequence element being unknown, to newly learned, and finally to being fully known. In the next analysis we tested the hypothesis of different functions in memory formation theorized for P300 and BP using finer temporal resolution. The P300 and BP progression of young (A) and older (B) participants are visualized on Figure 7A-B, which shows mean and standard error for P300 and BP amplitudes in each repetition relative to the point of first accurate recall. Both the P300 and BP amplitudes increase initially in young and remain unchanged in older up to the point of committing the stimulus to memory and then decrease gradually showing no distinct progressions.

**Figure 7.**
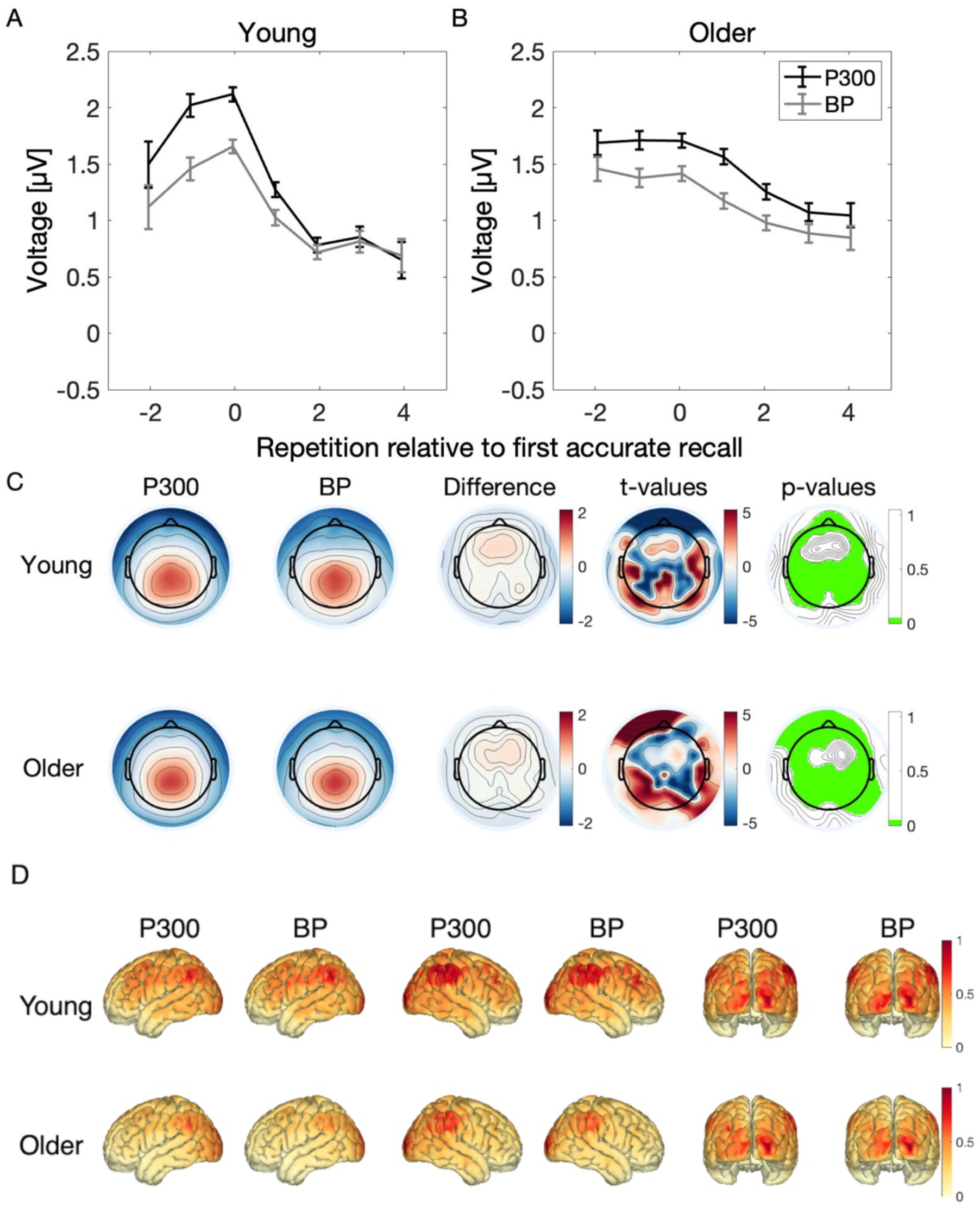
Distinct function of P300 and BP. (A, B) Gradual changes over the course of learning. In order to test the distinct functions in memory formation theorized for P300 and BP, we plotted the development of P300 (black) and BP (gray) amplitudes in young (A) and older (B) participants in relation to the point of first accurate recall (i.e newly learned; 0). Both P300 and BP amplitudes increase at first towards the point of first accurate recall and decrease afterwards. The gradual changes over the course of learning are almost equivalent for both components. Please note that different time points (i.e., repetitions relative to the newly learned point) can contain different sets of participants, because participants that learned all stimuli in the first repetition do not have any data in the time points -1 or -2. Error bars represent the standard error of the mean. (C) Topographical maps of P300, broad positivity, difference of both, and the results of equivalence tests computed on each electrode in young (top) and older (bottom) participants. The green color indicates electrodes with an equivalent activation between P300 and BP (i.e., significant equivalence test result), which means that for these electrodes one can reject the hypothesis that the true effect is smaller than d = -0.5 or larger than d = 0.5. In young as well as in older participants, the TOST procedure revealed equivalent centro-parietal topography for P300 and BP. (D) Source reconstructed spatial distribution of P300 and BP in young and older participants (left, right, posterior).

#### 3. 2. 7. Neural generators of P300 and BP

Since the P300 and BP are both centro-parietal positivities, and when measured relative to the same baseline exhibit similar changes during learning, we next sought to statistically test the hypothesis that they are generated in the same brain area. First, we investigate the differences in the distribution of neural activity in P300 and BP by performing a topographic analysis of variance (TANOVA). In young and older participants, the difference in scalp topographies was not statistically significant (DISS = 0.2158, p = 1 and DISS = 0.1553, p = 1, respectively). Figure 7C depicts the topography of P300, BP and the topographical difference of both components. The results of TANOVA indicate that the topographical maps of P300 and BP are not different, meaning that it is very unlikely that they are driven by different neural generators.

Next, to substantiate that the topographical maps of P300 and BP are equivalent, we computed a two one-sided equivalence test (TOST) on each electrode with the bound set to a medium effect size of d = 0.5 (Figure 7C). The results of the TOST procedure performed on all 105 electrodes confirmed mostly equivalent centro-parietal topography (green) for P300 and BP in young and older participants. The non-significant frontal regions represent residual activity, which resembles the next closest dominant component, the P3a. For a TOST with the bound set to the smallest effect size of interest (SESOI, (Lakens, 2017)) based on power analysis see Supplementary Figure 4.

Finally, we performed a source reconstruction analysis of P300 and BP to further examine the neural generators of both components. The results are visualized in Figure 7D. We observed a very similar distribution of neural generators of P300 and BP within young and older participants. The source reconstruction analysis revealed distributed activation patterns over fronto-parietal networks with the highest activation in the occipital and parietal brain regions. These results suggest that P300 and BP are indeed generated by the same brain regions.

### 3. 3. Reliability of behavioral performance and P300 amplitude

Finally, we computed the reliability of behavioral and electrophysiological measures related to learning to provide information on their within- and between sessions stability. Within the first session, we found fair reliability for the time constant tau (ICC = 0.57) and good reliability for P300 amplitude (ICC = 0.60). Within the second session, we found fair reliability for the time constant tau (ICC = 0.45) and fair reliability for P300 amplitude (ICC = 0.40). Furthermore, time constant tau showed excellent (ICC = 0.79) and P300 amplitude good (ICC = 0.67) test-retest reliability across one week (i.e., two sessions). Sensitivity analysis revealed that the ICCs were robust across both age groups. Our results demonstrate that the time constant tau provides fair to excellent reliability and P300 amplitude provides fair to good reliability to measure learning performance and neurophysiological processes related to learning and memory formation.

## Discussion

In the present study, we investigated the effects of aging on learning trajectories and memory formation. By analyzing behavioral and neurophysiological indices during a visuospatial sequence learning task, we were able to gain insights into the neurophysiological mechanisms of gradual memory formation in young and older healthy participants. The employed sequence learning paradigm, in which a fixed sequence of spatially distinct stimuli was memorized over repeated observations, enables to track the gradual nature of learning that are precluded in traditional approaches utilizing a remembered vs. forgotten comparison.

On a behavioral level, we first demonstrated that both young and older participants were able to learn the sequence and improved with each sequence repetition. However, the analysis of the *learning index* (LI) revealed that young participants learned significantly faster compared to older participants and the younger participants memorized more sequence elements than older participants, which was illustrated by the substantial age-effect of the *knowledge index* (KI), which is in line with previous memory aging research (Fernandes & Wammes, 2015; Nyberg et al., 2012; Riddle, 2007; Werkle-Bergner et al., 2006).

The analysis of the neurophysiological data revealed a continuous decrease of the P300 as stimulus locations were better learned and hence increasingly anticipated. The decrease of the P300 was larger in young participants, which was significantly linked to an increasing knowledge about the learned material (i.e., KI). Crucially, our study showed that within-subject variation of learning was significantly associated with the decrease of the P300 amplitude. In line with this, a subsequent analysis demonstrated, consistently across both age groups, that the P300 and BP amplitudes varied systematically across the specific learning states (i.e., unknown, newly learned, known) during the sequence memorization. In both age groups, the two ERP measures did not significantly change from unknown to newly learned states, but strongly decreased after committing the stimuli to memory. However, between age group comparisons exhibited in young subjects significantly greater P300 amplitude in newly learned and significantly smaller amplitude in known stimuli. Moreover, we demonstrated that the behavioral performance and P300 amplitude are reliable measures within and between sessions. Taken together, the results show that the less effective sequence memorisation evident in the behavioral reports of older participants, is systematically reflected in the more gradual decrease over repetitions of neural activity components reflecting the degree of expectancy and the process of new memory formation, highlighting promise in tracking learning effectiveness independently of behavioral reporting abilities. In the following we incorporate the findings into the current theories about centro-parietal ERP components and memory formation in older participants.

### The role of centroparietal ERP activity during memory formation

Previous ERP research on memory revealed increased broad centro-parietal positivity approximately 300-800 ms after stimulus for items later being remembered compared to those forgotten in various tasks (Eimer et al., 1996; Gonsalves & Paller, 2000; Neville et al., 1986; Paller et al., 1987, 1988; Schlaghecken et al., 2000). This “difference in subsequent memory” effect has been observed in a wide range of encoding conditions, although other studies failed to find this effect (for a review see: (Johnson, 1995)). However, some studies argued that centro-parietal amplitude is not primarily associated with the memorization itself, but rather mediated by stimulus complexity and semantic processing depth (Petten & Senkfor, 1996). Using simple visual stimuli with minimal semantic value as in the present sequence learning task, where there are no differences in complexity across items, we provide evidence that these modulations of centro-parietal ERP activity are related to memory formation. This is further substantiated by the significant differences between different learning states (i.e., unknown, newly learned and known). Although stimuli of low semantic content have been used previously, the majority of these studies analyzed the neurophysiological signature during the recognition rather than encoding (Curran, 2004; Curran & Cleary, 2003; Gonsalves & Paller, 2000; Voss & Paller, 2009). As the recognition process is inseparably compounded with decision processes to determine if the stimulus was seen before (Curran, 2004; Voss & Paller, 2009), the exact role of the P300 during encoding remained unclear. Moreover, these traditional memory examination methods had the limitation that their design did not allow to track the gradual memory formation process. We addressed this specifically in our study by utilizing a paradigm which naturally provides the opportunity to investigate the incremental nature of learning. Therefore, we were able not only to compare the P300/BP amplitude of remembered and not remembered stimuli, but examine how the amplitude changes during learning, from stimulus being unknown, to newly learned and finally fully known. An additional motivation for the distinction of these different learning states is that the relationship between the modulations of P300, BP and over the course of learning may simply reflect the habituation processes (Polich, 1989; Ravden & Polich, 1998). Therefore, we aimed to provide a stronger test for the relationship between neural signals and memory formation by categorizing the stimuli into different learning states: unknown, newly learned and known and examined amplitude modulations independently of the sequence repetition number.

Consistently across age groups, contrast comparisons demonstrated that both components showed significantly greater amplitude for unknown and newly learned stimuli compared to known stimuli. The difference between unknown and newly learned stimuli did not reach significance. When comparing both age groups, young had significantly greater P300 amplitude for newly learned and significantly smaller amplitude for known stimuli. However, contrast comparisons showed no age differences in BP amplitude across learning categories. Interestingly, we found no age-related difference in P300 amplitude of unknown stimuli. Although, when examining the P300 amplitude over sequence repetitions, young exhibit greater amplitude in the first repetition and a steeper monotonic decrease afterwards (Figure 5B). The fact that unknown P300 amplitudes were not elevated in the young participants despite elevated P300s in repetition #1 might be explained by the fact that a large portion of the young participants learned the majority of the sequence elements already during the first sequence repetition. The unknown category therefore consists of the subset of sequences and participants for which learning was slower, and to the extent that this was linked with lower attentional resources, it may be associated with relatively smaller P300 amplitudes (Polich, 2007). Indeed, the faster learning rate in young participants might be in some part attributable to the relatively large amplitude and short latency of the P300 peak for newly learned stimuli, as more efficient stimulus identification likely facilitates better memorisation of those stimuli. Further, the drop of the P300 amplitude after committing the stimulus to memory (i.e., from newly learned to known) was steeper in young than in older participants, as shown by the significant interaction of age group and categories newly learned (i.e., intercept) and known. Moreover, young exhibit smaller P300 amplitudes of the known stimuli compared to older participants. These results confirm the construct of uncertainty to be reflected by P300 and may suggest that in this undemanding task young people exhibit high levels of confidence in their knowledge shortly after committing the stimulus to memory, whereas older participants still express some level of doubt.

One obvious prediction could have been that older people are less efficient learners because their memory activity is weaker. However, we don’t see evidence for such an overall effect. The lack of appreciable reduction in BP amplitude in the old compared to young is remarkable. It seems that despite the memory formation process being engaged to a similar strength, this engagement does not translate into the same learning progress in the older participants. One possibility is that the older participants employ the memory formation process to a similar overall extent but less strategically, e.g., trying equally hard to memorize all stimuli at once rather than break the sequence down into chunks. Alternatively, it could be that the slower and more variable timing of the stimulus identification process reflected in the P300 means that despite the older participants engaging the memory formation process, there is less time for it to translate the categorical stimulus location information into a solidified memory trace. Whatever the underlying reason, the fact that the BP is less effective at generating strong memory traces is borne out in our analysis showing lower beta weights linking the BP to learning rate in the older participants.

A core principle in this study was that P300 amplitude decreases as a function of subjective stimulus expectancy (Donchin, 1981; Duncan-Johnson & Donchin, 1977; Kolossa et al., 2012; Mars et al., 2008; Rüsseler et al., 2003; Schlaghecken et al., 2000; Sutton et al., 1965), which in the context of sequence learning corresponds to knowledge about the sequence. However, a recent study advocated for an attentional role of the P300, such that it should increase for stimuli as they are better stored in memory, which is opposite to the expectancy/surprise account (Jongsma et al., 2012). We found that P300 decreased over the course of learning, which is in line with the decremental surprise linked to increasingly confident sequence knowledge. Expectancy-driven P300 modulations have been also linked to working memory processes (Karis et al., 1984; Polich, 2007; Steiner et al., 2013; Verleger, 2020). This suggests that the link between expectancy and learning may extend beyond the sequence memorization task studied here. From an evolutionary point of view, rare events may be extremely important, therefore should be stored more intensely in the working memory than events that occur frequently. Additionally, in order to correctly respond to an event later, one must be held in working memory first. Consistent with previous research, we demonstrated lower performance and delayed P300 peak latencies (Fjell & Walhovd, 2001; Polich, 1997; Walhovd et al., 2008). Lower performance of older people may be linked to previously reported slowing processing speed, which might be caused by decreased axon myelination and reduction in neurotransmitter levels (Park & Festini, 2017; Van Petten et al., 2004), and the longer P300 peak latencies may reflect the time spent on stimulus categorization (Teixeira-Santos et al., 2020). The fact that older participants exhibited a delayed P300 latency might indicate that older participants required more time for stimulus evaluation. Moreover, the delayed peak latencies may damage the memory formation process because they leave less time to integrate the current stimulus location with previous or next position in the sequence.

### Distinct functions of P300 and BP?

The modulations of P300 and BP amplitude across repetitions and learning states were highly similar, which may question the assumption of two distinct components reflecting different processes in learning. Indeed, whether BP is a distinct component or just a prolonged P300 is still open to debate (Hajcak & Foti, 2020; Verleger, 2020). Previously, (Steinemann et al., 2016) attempted to isolate components within the P300 wave by conducting a finer temporal analysis and examining the development of both neural signals in relation to the point of first accurate recall (i.e., newly learned). Their rationale was that two overlapping components with different timescales can be measured by locally baselining the faster one (i.e., measuring P300 peak relative to a preceding trough) and selecting a time window beyond the faster component to measure the broader one. Using this method, Steineman et al. (2016) was able to demonstrate different progressions of P300 and BP during learning, where the BP did not begin to drop until after the point at which an item became newly learned, whereas the P300 dropped monotonically starting from the first repetition even before becoming known, reflecting a more gradual change in expectancy. This supported their contention that the P300 represents expectancy and surprise encoding whereas the function of the more sustained BP activity is to hold previous and future items in working memory in order to facilitate active memory formation by forming sequential relations with the current item. The key difference in the current study was that the P300, like the BP, did not begin decreasing before the item became newly learned. A potential explanation is the difference in baselining methods - here, a local baseline (i.e., trough-to-peak) for the P300 proved problematic because the waveforms did not suggest a very clear preceding trough, and indeed this lack of clear guidance for baseline interval definition is a general shortcoming of the local-baselining approach. Without local baselining, the transient P300 may pick up the same modulations as the BP due to being superimposed on it. However, this still does not fully explain the discrepancy, because a supplementary analysis using trough-to-peak measurement for the P300 also did not replicate the monotonic P300 reduction in advance of memory acquisition (Supplementary Figure 3). When aligned to the point of first accurate recall, the P300 amplitude still increased (in young) or remained stable (in older) up to this point, followed by a continuous decrease for already known stimuli. A potential alternative explanation lies in the difference in difficulty of the current version of the task, where young participants were able to recall more than 65% of the sequence after one repetition, in contrast to the previous study where subjects could recall only approximately 50% even after three repetitions. This means that there was a much smaller proportion of the trials in the young that were categorized as unknown, and it could be that these fewer instances of failed immediate learning were more closely linked with decreased attention or weaker stimulus identification, bringing their amplitude down close to the level of all stimuli when newly learned. Taken together, the current results cast doubt on the generality of the differences in learning trajectories found for the early (P300) and broader (BP) in Steinemann et al (2016) and underline the need for more research into whether and how they dissociate from each other in the context of learning. The gradual decrease of amplitude over the course of several repetitions after committing the stimuli to memory might reflect the increasing confidence about the sequence, as correct recall might not be synonymous with having no doubts.

In addition to the similar learning trajectories, the P300 and BP exhibited similar topographies, raising the question of whether they are generated in the same brain regions. First, we compared the topographical maps of both components using TANOVA and equivalence tests on sensor level. The idea behind those tests was based on the premise that it is very unlikely that different neural generators produced the same scalp topography (Koenig et al., 2011; Murray et al., 2008). The results of TANOVA showed no differences between topographies of P300 and BP in young and older participants. These results indicate that both components do not reflect distinct processes during learning. Similarly, the results of equivalence tests showed equivalent posterior topography of P300 and BP indicating the same sources of activation. The topographical maps of P300 and BP were not equivalent on a set of frontal electrodes, however this residual frontal activity might represent the next dominant component, which is probably the frontal P3a. Finally, the source reconstruction analysis revealed concordant distributed brain activation patterns within parietal circuits with the highest activation in the occipital and parietal brain regions. Taken together, in our sample and paradigm, there is no evidence (rather evidence for equivalence) for distinct underlying neural generators for the two components.

### The P300 as a biomarker to predict learning success

Having established the evidence for the contribution of the P300 in sequence learning, we subsequently investigated if these components were effective in predicting learning success within and across participants and potentially could serve as a biomarker for successful learning. To qualify as a biomarker, a prerequisite is a sufficient reliability of these neurophysiological and behavioral measures. Critically, our results demonstrated fair to excellent within and between sessions reliability for the behavioral performance and the P300 amplitude. The high within-subject reliability fulfills a vital criterion for a potential useful biomarker.

Next, we examined the within-subject prediction by testing if the P300 and broad positivity were predictive for the behavioral measures defined as the knowledge index and learning index. In both age groups, the results revealed that the average P300 amplitude in a given repetition could predict the cumulative knowledge about the sequence (i.e., knowledge index) within that repetition. In addition, there was a positive association between broad positivity and learning rate (i.e., learning index). The significant interaction of both neural signals with the age group indicated that the association was stronger in young compared to older participants. We further demonstrated that inter-individual learning success can be predicted by the P300 amplitude among young, but not older individuals. Specifically, the decrease of P300 amplitude in repetitions, where most of the learning occurred, was able to identify the fastest learners in young participants. A possible explanation for the less reliable prediction of learning success in older participants might be changing functional organisation of the brain, which becomes less organized and more random with increasing age (Petti et al., 2016). (Knyazev et al., 2015) reached a similar conclusion, by demonstrating that the older participants exhibit lower connectivity in beta and gamma band networks. Further, in some studies older participants show increased frontal P300, which might reflect compensatory or inefficient use of neural resources (Fjell & Walhovd, 2001; Saliasi et al., 2013). The reorganization and increasingly random activation of the brain may make reliable predictions of learning more difficult in older age. Another cause of poor recall in the older people could be that despite similar to young engagement of memory formation processes not all of the learned material translated efficiently into solid sequence knowledge possibly due to poor learning strategy, or even if they are learning, they might make more mistakes in reporting that learning. The older people may suffer interference effects during the recall itself while spending resources on using computer mouse which compromise their very recently-made, still precarious memory trace and so they lose it. The younger people’s memory traces would be more resilient to having to use a mouse to report the locations, because they are more familiar with technology.

### Limitations

One limitation of the present study might be the low task difficulty for young participants. Many young participants learned the sequence of stimuli already after the initial sequence presentation leading to no (or only few) UN, but only NL and K stimuli. This resulted in an unbalanced design with the majority of K stimuli and only a small portion of UN stimuli. However, we adjusted the task difficulty in a pilot study prior to data collection to the capabilities of the older participants, as the main goal of this study was to investigate the age-related learning trajectories. Moreover, we used linear mixed-effect models for statistical analysis to account for an unbalanced number of stimuli in each learning category.

Finally, we did not account for the learning strategy used by the participants. As mentioned above, the P300 is not elicited, when participants chose to ignore certain stimuli in a given sequence repetition in order to learn them in the next one. Therefore, the low amplitudes in the ‘-2’ and ‘-1’ categories (i.e., two and one repetition prior to the point of first accurate recall, respectively) might be the result of not engaging attentional resources, and not expectancy or learning. Therefore, we also examined the amplitude modulations across sequence repetitions, where the effects of learning strategy play a less important role.

## Conclusion

Using a paradigm that provides the opportunity to investigate the incremental nature of learning, we gained novel insights into aging effects of neural mechanisms underlying the gradual memory formation process. Our data demonstrate that the diminished learning capabilities in older participants were directly linked to more gradual modulations of the P300 and BP amplitude over the course of learning reflecting more spread-out learning progress. The decreased amplitude may be interpreted as evidence of cognitive decline caused by deterioration in posterior brain regions and functional reorganization of the brain, which mean that older participants require more time for stimulus evaluation and engage a lower level of attentional resources resulting in lower behavioral performance and less distinct confidence about the learned material. We provide evidence that the P300 amplitude can predict learning success across sequence repetitions in both age groups. The present report foregrounds the important role of the expectancy-driven P300 as potential biomarker for learning success, which may facilitate the development of preventive techniques for age-related impeded learning.

## Acknowledgment

This work was supported by the Swiss National Science Foundation.

## Data Availability

All data, preprocessing and analysis scripts used for the analyses are uploaded on OSF.io at https://osf.io/sgnmx/. We agree to share our data, any digital study materials, and laboratory logs for all published results in this repository.

## Abbreviations

EEG: electroencephalography
ERP: event-related potential
BP: broad positivity
mP300: average P300
mBP: average broad positivity
MMSE: mini-mental state exam
AD: Alzheimer’s disease
UN: unknown
NL: newly learned
K: known
F: forgotten
KI: knowledge index
LI: learning index
ICA: independent component analysis
TANOVA: topographic analysis of variance
DISS: dissimilarity index
TOST: two one-sided equivalence tests
ICC: intraclass correlation coefficient

## References

Alday, P. M. (2019). How much baseline correction do we need in ERP research? Extended GLM model can replace baseline correction while lifting its limits. Psychophysiology, 56(12), e13451.

Bigdely-Shamlo, N., Mullen, T., Kothe, C., Su, K.-M., & Robbins, K. A. (2015). The PREP pipeline: standardized preprocessing for large-scale EEG analysis. Frontiers in Neuroinformatics, 9, 16.

Chiang, H.-S., Hsiao, K.-L., & Liu, L.-C. (2018). EEG-Based Detection Model for Evaluating and Improving Learning Attention. Journal of Medical and Biological Engineering, 38(6), 847– 856.

Cicchetti, D. V. (1994). Guidelines, criteria, and rules of thumb for evaluating normed and standardized assessment instruments in psychology. Psychological Assessment, 6(4), 284– 290.

Curran, T. (2004). Effects of attention and confidence on the hypothesized ERP correlates of recollection and familiarity. Neuropsychologia, 42(8), 1088–1106.

Curran, T., & Cleary, A. M. (2003). Using ERPs to dissociate recollection from familiarity in picture recognition. Brain Research. Cognitive Brain Research, 15(2), 191–205.

DeCarli, C. (2003). Mild cognitive impairment: prevalence, prognosis, aetiology, and treatment. Lancet Neurology, 2(1), 15–21.

de Cheveigné, A. (2020). ZapLine: A simple and effective method to remove power line artifacts. NeuroImage, 207, 116356.

Delorme, A., & Makeig, S. (2004). EEGLAB: an open source toolbox for analysis of single-trial EEG dynamics including independent component analysis. Journal of Neuroscience Methods, 134(1), 9–21.

Donchin, E. (1981). Surprise!? Surprise? Psychophysiology, 18(5), 493–513.

Duncan-Johnson, C. C., & Donchin, E. (1977). On quantifying surprise: the variation of event-related potentials with subjective probability. Psychophysiology, 14(5), 456–467.

Eimer, M., Goschke, T., Schlaghecken, F., & Stürmer, B. (1996). Explicit and implicit learning of event sequences: evidence from event-related brain potentials. Journal of Experimental Psychology. Learning, Memory, and Cognition, 22(4), 970–987.

Fernandes, M. A., & Wammes, J. D. (2015). Memory and Memory Theory. In The Encyclopedia of Adulthood and Aging (pp. 1–5). https://doi.org/10.1002/9781118521373.wbeaa123

Fjell, A. M., & Walhovd, K. B. (2001). P300 and neuropsychological tests as measures of aging: scalp topography and cognitive changes. Brain Topography, 14(1), 25–40.

Folstein, M. F., Folstein, S. E., & McHugh, P. R. (1975). “Mini-mental state”: a practical method for grading the cognitive state of patients for the clinician. Journal of Psychiatric Research, 12(3), 189–198.

Frömer, R., Maier, M., & Abdel Rahman, R. (2018). Group-Level EEG-Processing Pipeline for Flexible Single Trial-Based Analyses Including Linear Mixed Models. Frontiers in Neuroscience, 12, 48.

Gonsalves, B., & Paller, K. A. (2000). Neural events that underlie remembering something that never happened. Nature Neuroscience, 3(12), 1316–1321.

Grady, C. L., Protzner, A. B., Kovacevic, N., Strother, S. C., Afshin-Pour, B., Wojtowicz, M., Anderson, J. A. E., Churchill, N., & McIntosh, A. R. (2010). A multivariate analysis of age-related differences in default mode and task-positive networks across multiple cognitive domains. Cerebral Cortex, 20(6), 1432–1447.

Gramfort, A., Papadopoulo, T., Olivi, E., & Clerc, M. (2010). OpenMEEG for M/EEG forward modeling: a comparison study. Human Brain Mapping. https://hal.inria.fr/inria-00502745/

Hajcak, G., & Foti, D. (2020). Significance?& Significance! Empirical, methodological, and theoretical connections between the late positive potential and P300 as neural responses to stimulus significance: An integrative review. Psychophysiology, 57(7), e13570.

Hämäläinen, M. S., & Ilmoniemi, R. J. (1994). Interpreting magnetic fields of the brain: minimum norm estimates. Medical & Biological Engineering & Computing, 32(1), 35–42.

Hansenne, M. (2000). The p300 cognitive event-related potential. II. Individual variability and clinical application in psychopathology. Neurophysiologie Clinique= Clinical Neurophysiology, 30(4), 211–231.

Harrison, X. A., Donaldson, L., Correa-Cano, M. E., Evans, J., Fisher, D. N., Goodwin, C. E. D., Robinson, B. S., Hodgson, D. J., & Inger, R. (2018). A brief introduction to mixed effects modelling and multi-model inference in ecology. PeerJ, 6, e4794.

Johnson, R. (1995). Event-related potential insights into the neurobiology of memory systems. Handbook of Neuropsychology, 10, 135–135.

Jongsma, M. L. A., Gerrits, N. J. H. M., van Rijn, C. M., Quiroga, R. Q., & Maes, J. H. R. (2012). Event related potentials to digit learning: tracking neurophysiologic changes accompanying recall performance. International Journal of Psychophysiology: Official Journal of the International Organization of Psychophysiology, 85(1), 41–48.

Karis, D., Fabiani, M., & Donchin, E. (1984). “P300” and memory: Individual differences in the von Restorff effect. Cognitive Psychology, 16(2), 177–216.

Knyazev, G. G., Volf, N. V., & Belousova, L. V. (2015). Age-related differences in electroencephalogram connectivity and network topology. Neurobiology of Aging, 36(5), 1849–1859.

Koenig, T., Kottlow, M., Stein, M., & Melie-García, L. (2011). Ragu: a free tool for the analysis of EEG and MEG event-related scalp field data using global randomization statistics. Computational Intelligence and Neuroscience, 2011, 938925.

Kolossa, A., Fingscheidt, T., Wessel, K., & Kopp, B. (2012). A model-based approach to trial-by-trial p300 amplitude fluctuations. Frontiers in Human Neuroscience, 6, 359.

Kuznetsova, A., Brockhoff, P. B., & Christensen, R. H. B. (2017). lmerTest Package: Tests in Linear Mixed Effects Models. *Journal of Statistical Software*, Articles, 82(13), 1–26.

Lakens, D. (2017). TOSTER: Two one-sided tests (TOST) equivalence testing. R Package Version 0. 2, 5, 648.

Langer, N., Ho, E. J., Alexander, L. M., Xu, H. Y., Jozanovic, R. K., Henin, S., Petroni, A., Cohen, S., Marcelle, E. T., Parra, L. C., Milham, M. P., & Kelly, S. P. (2017). A resource for assessing information processing in the developing brain using EEG and eye tracking. Scientific Data, 4, 170040.

Lenth, R. (2020). Emmeans: Estimated Marginal Means, aka Least-Squares Means, R package version 1.*4*. 5.; 2020.

Liesefeld, H. R. (2018). Estimating the Timing of Cognitive Operations With MEG/EEG Latency Measures: A Primer, a Brief Tutorial, and an Implementation of Various Methods. Frontiers in Neuroscience, 12, 765.

Luu, P., Poulsen, C., & Tucker, D. M. (2009). Neurophysiological Measures of Brain Activity: Going from the Scalp to the Brain. Foundations of Augmented Cognition. Neuroergonomics and Operational Neuroscience, 488–494.

Mars, R. B., Debener, S., Gladwin, T. E., Harrison, L. M., Haggard, P., Rothwell, J. C., & Bestmann, S. (2008). Trial-by-trial fluctuations in the event-related electroencephalogram reflect dynamic changes in the degree of surprise. The Journal of Neuroscience: The Official Journal of the Society for Neuroscience, 28(47), 12539–12545.

McGraw, K. O., & Wong, S. P. (1996). Forming inferences about some intraclass correlation coefficients. Psychological Methods, 1(1), 30–46.

Moisello, C., Meziane, H. B., Kelly, S., Perfetti, B., Kvint, S., Voutsinas, N., Blanco, D., Quartarone, A., Tononi, G., & Ghilardi, M. F. (2013). Neural activations during visual sequence learning leave a trace in post-training spontaneous EEG. PloS One, 8(6), e65882.

Murray, M. M., Brunet, D., & Michel, C. M. (2008). Topographic ERP analyses: a step-by-step tutorial review. Brain Topography, 20(4), 249–264.

Neville, H. J., Kutas, M., Chesney, G., & Schmidt, A. L. (1986). Event-related brain potentials during initial encoding and recognition memory of congruous and incongruous words. Journal of Memory and Language, 25(1), 75–92.

Nyberg, L., Lövdén, M., Riklund, K., Lindenberger, U., & Bäckman, L. (2012). Memory aging and brain maintenance. In Trends in Cognitive Sciences (Vol. 16, Issue 5, pp. 292–305). https://doi.org/10.1016/j.tics.2012.04.005

Olkin, I. (1960). Contributions to probability and statistics: essays in honor of Harold Hotelling. Stanford University Press.

Paller, K. A., Kutas, M., & Mayes, A. R. (1987). Neural correlates of encoding in an incidental learning paradigm. Electroencephalography and Clinical Neurophysiology, 67(4), 360–371.

Paller, K. A., McCarthy, G., & Wood, C. C. (1988). ERPs predictive of subsequent recall and recognition performance. Biological Psychology, 26(1-3), 269–276.

Park, D. C., & Festini, S. B. (2017). Theories of memory and aging: A look at the past and a glimpse of the future. The Journals of Gerontology. Series B, Psychological Sciences and Social Sciences. https://academic.oup.com/psychsocgerontology/article-pdf/doi/10.1093/geronb/gbw066/13798644/gbw066.pdf

Patterson, J. V., Michalewski, H. J., & Starr, A. (1988). Latency variability of the components of auditory event-related potentials to infrequent stimuli in aging, Alzheimer-type dementia, and depression. Electroencephalography and Clinical Neurophysiology, 71(6), 450–460.

Pedroni, A., Bahreini, A., & Langer, N. (2019). Automagic: Standardized preprocessing of big EEG data. NeuroImage, 200, 460–473.

Pernet, C. R., Chauveau, N., Gaspar, C., & Rousselet, G. A. (2011). LIMO EEG: a toolbox for hierarchical LInear MOdeling of ElectroEncephaloGraphic data. Computational Intelligence and Neuroscience, 2011, 831409.

Petten, C., & Senkfor, A. J. (1996). Memory for words and novel visual patterns: Repetition, recognition, and encoding effects in the event-related brain potential. In Psychophysiology (Vol. 33, Issue 5, pp. 491–506). https://doi.org/10.1111/j.1469-8986.1996.tb02425.x

Petti, M., Toppi, J., Babiloni, F., Cincotti, F., Mattia, D., & Astolfi, L. (2016). EEG Resting-State Brain Topological Reorganization as a Function of Age. Computational Intelligence and Neuroscience, 2016, 6243694.

Pion-Tonachini, L., Kreutz-Delgado, K., & Makeig, S. (2019). ICLabel: An automated electroencephalographic independent component classifier, dataset, and website. NeuroImage, 198, 181–197.

Polich, J. (1989). Habituation of P300 from auditory stimuli. Psychobiology, 17(1), 19–28.

Polich, J. (1997). On the relationship between EEG and P300: individual differences, aging, and ultradian rhythms. International Journal of Psychophysiology: Official Journal of the International Organization of Psychophysiology, 26(1-3), 299–317.

Polich, J. (2007). Updating P300: an integrative theory of P3a and P3b. Clinical Neurophysiology: Official Journal of the International Federation of Clinical Neurophysiology, 118(10), 2128– 2148.

Polich, J., Howard, L., & Starr, A. (1985). Effects of Age on the P300 Component of the Event-related Potential From Auditory Stimuli: Peak Definition, Variation, and Measurement1. Journal of Gerontology, 40(6), 721–726.

Porcaro, C., Balsters, J. H., Mantini, D., Robertson, I. H., & Wenderoth, N. (2019). P3b amplitude as a signature of cognitive decline in the older population: An EEG study enhanced by Functional Source Separation. NeuroImage, 184, 535–546.

Ravden, D., & Polich, J. (1998). Habituation of P300 from visual stimuli. International Journal of Psychophysiology: Official Journal of the International Organization of Psychophysiology, 30(3), 359–365.

Riddle, D. R. (2007). Brain Aging: Models, Methods, and Mechanisms. CRC Press.

Rüsseler, J., Hennighausen, E., Münte, T. F., & Rösler, F. (2003). Differences in incidental and intentional learning of sensorimotor sequences as revealed by event-related brain potentials. Brain Research. Cognitive Brain Research, 15(2), 116–126.

Saliasi, E., Geerligs, L., Lorist, M. M., & Maurits, N. M. (2013). The relationship between P3 amplitude and working memory performance differs in young and older adults. PloS One, 8(5), e63701.

Schlaghecken, F., Stürmer, B., & Eimer, M. (2000). Chunking processes in the learning of event sequences: electrophysiological indicators. Memory & Cognition, 28(5), 821–831.

Steinemann, N. A., Moisello, C., Ghilardi, M. F., & Kelly, S. P. (2016). Tracking neural correlates of successful learning over repeated sequence observations. NeuroImage, 137, 152–164.

Steiner, G. Z., Barry, R. J., & Gonsalvez, C. J. (2013). Can working memory predict target-to-target interval effects in the P300? International Journal of Psychophysiology: Official Journal of the International Organization of Psychophysiology, 89(3), 399–408.

Sutton, S., Braren, M., Zubin, J., & John, E. R. (1965). Evoked-potential correlates of stimulus uncertainty. Science, 150(3700), 1187–1188.

Tanner, D., Morgan-Short, K., & Luck, S. J. (2015). How inappropriate high-pass filters can produce artifactual effects and incorrect conclusions in ERP studies of language and cognition. Psychophysiology, 52(8), 997–1009.

Teixeira-Santos, A. C., Pinal, D., Pereira, D. R., Leite, J., Carvalho, S., & Sampaio, A. (2020). Probing the relationship between late endogenous ERP components with fluid intelligence in healthy older adults. Scientific Reports, 10(1), 11167.

Tinga, A. M., de Back, T. T., & Louwerse, M. M. (2019). Non-invasive neurophysiological measures of learning: A meta-analysis. Neuroscience and Biobehavioral Reviews, 99, 59–89.

Tröndle, M., Popov, T., & Langer, N. (2020). Decomposing the role of alpha oscillations during brain maturation. In bioRxiv (p. 2020.11.06.370882). https://doi.org/10.1101/2020.11.06.370882

Tröndle, M., Popov, T., Pedroni, A., Pfeiffer, C., Barańczuk-Turska, Z., & Langer, N. (2021). Decomposing age effects in EEG alpha power. In bioRxiv (p. 2021.05.26.445765). https://doi.org/10.1101/2021.05.26.445765

van Deursen, J. A., Vuurman, E. F. P. M., Smits, L. L., Verhey, F. R. J., & Riedel, W. J. (2009). Response speed, contingent negative variation and P300 in Alzheimer’s disease and MCI. Brain and Cognition, 69(3), 592–599.

Van Petten, C., Plante, E., Davidson, P. S. R., Kuo, T. Y., Bajuscak, L., & Glisky, E. L. (2004). Memory and executive function in older adults: relationships with temporal and prefrontal gray matter volumes and white matter hyperintensities. Neuropsychologia, 42(10), 1313– 1335.

Verleger, R. (2020). Effects of relevance and response frequency on P3b amplitudes: Review of findings and comparison of hypotheses about the process reflected by P3b. Psychophysiology, 57(7), e13542.

Voss, J. L., & Paller, K. A. (2009). An electrophysiological signature of unconscious recognition memory. Nature Neuroscience, 12(3), 349–355.

Walhovd, K. B., Rosquist, H., & Fjell, A. M. (2008). P300 amplitude age reductions are not caused by latency jitter. Psychophysiology, 45(4), 545–553.

Werkle-Bergner, M., Müller, V., Li, S.-C., & Lindenberger, U. (2006). Cortical EEG correlates of successful memory encoding: implications for lifespan comparisons. Neuroscience and Biobehavioral Reviews, 30(6), 839–854.

Widmann, A., & Schröger, E. (2012). Filter effects and filter artifacts in the analysis of electrophysiological data. Frontiers in Psychology, 3, 233.

Wilkinson, G. N., & Rogers, C. E. (1973). Symbolic description of factorial models for analysis of variance. Journal of the Royal Statistical Society. Series C, Applied Statistics, 22(3), 392.

