## Supplementary Material for "Neurophysiological markers of successful learning in healthy aging"

### Supplementary Figures:

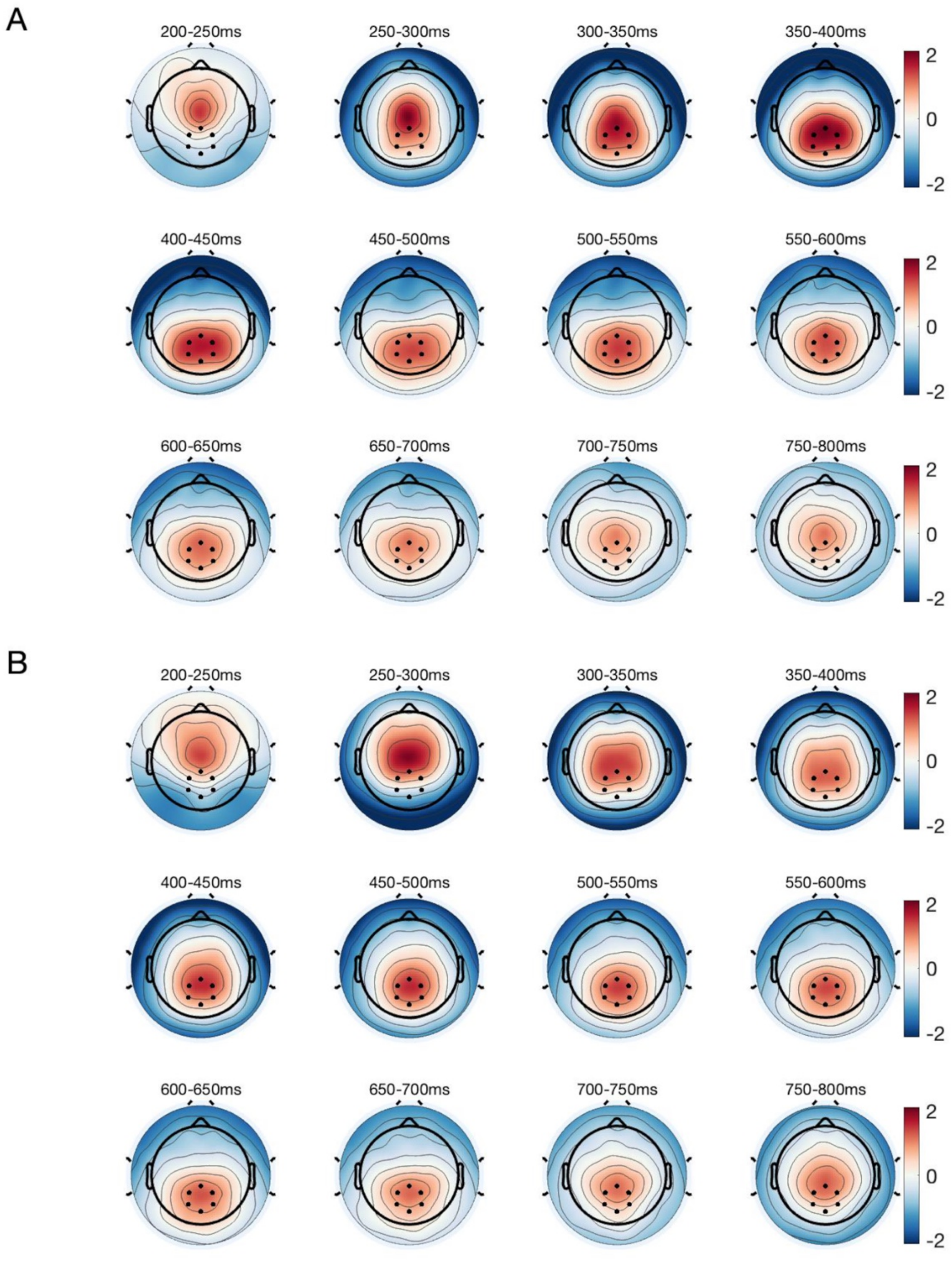


Supplementary Figure 1. Grand average scalp topographies plotted with a 50 ms step in young (A) and older (B) participants. The Grand average consists only of unknown and newly learned trials. The black dots indicate centro-parietal electrodes selected for computing ERPs and further statistical analysis.

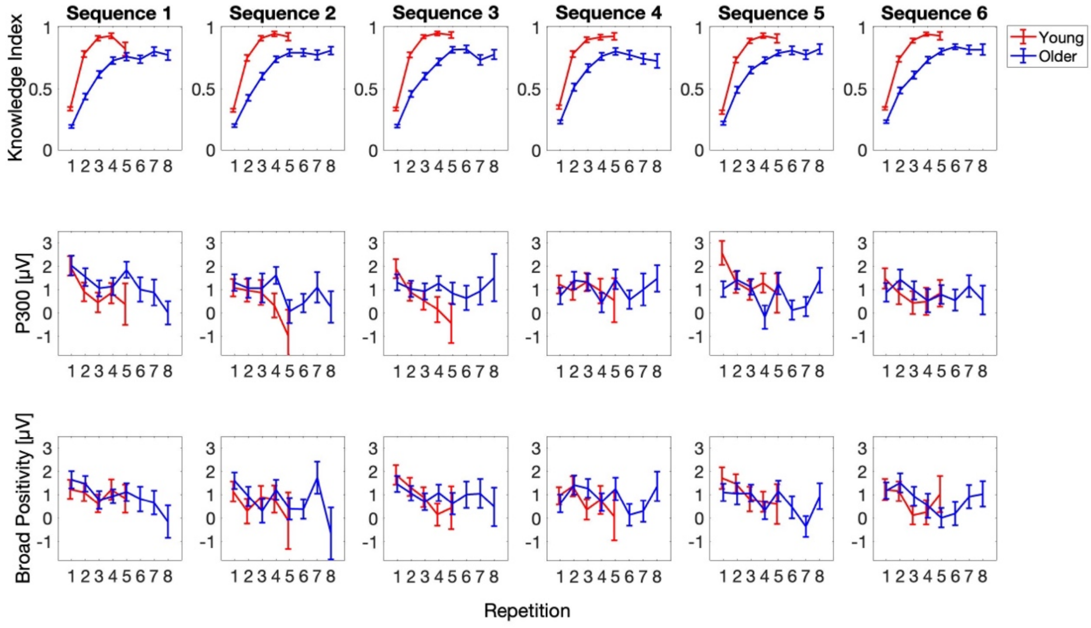


Supplementary Figure 2. Knowledge index, P300 and broad positivity across all six stimulus sequences (three from the first and three from the second session) in young (red) and older (blue) participants. The average knowledge index increased monotonically in young and older individuals; however, the young learned the sequence of stimuli faster than the older. Moreover, P300 and BP amplitudes decreased on average with increasing knowledge index in both age groups. In young, only the first five sequence repetitions were plotted due to the low number of trials from the sixth repetition on. Error bars represent the standard error of the mean.

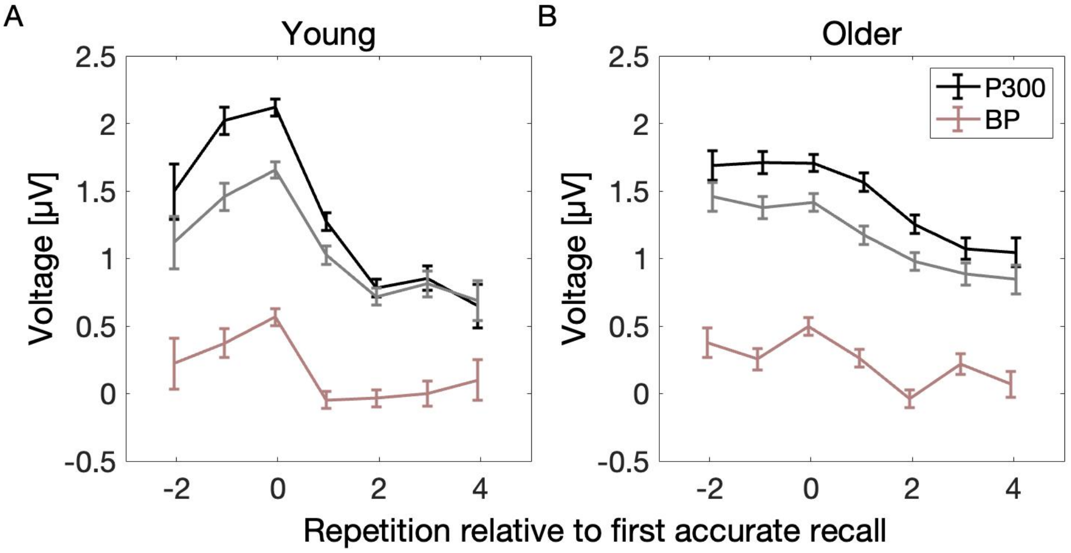


Supplementary Figure 3. Gradual changes over the course of learning. In order to test the distinct functions in memory formation theorized for P300 and BP, we plotted the development of P300 (black), P300 trough-to-peak (TTP; beige) and BP (gray) amplitudes in young (left) and older (right) participants in relation to the point of first accurate recall (i.e., newly learned; 0). P300, TTP and BP amplitudes increase at first towards the point of first accurate recall and decrease afterwards. The gradual changes over the course of learning are almost equivalent for all components. Error bars represent the standard error of the mean.

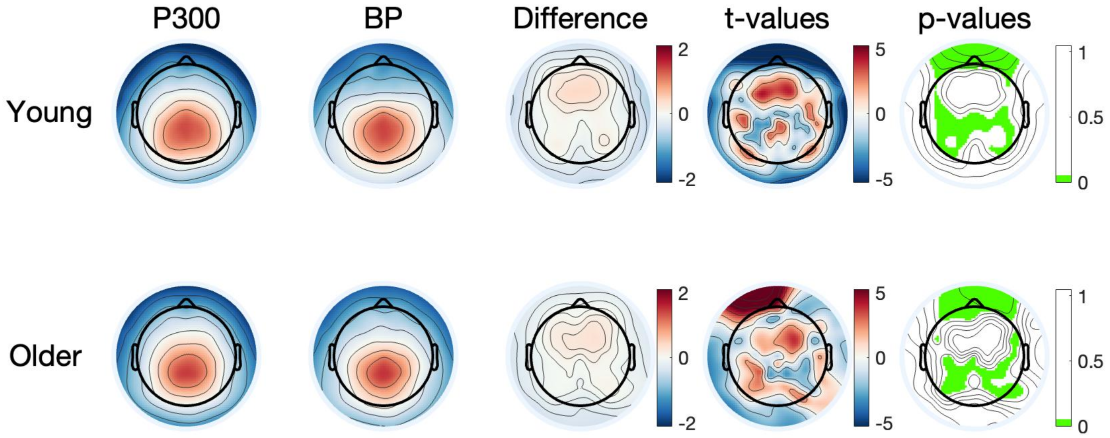


Supplementary Figure 4. Topographical maps of P300, broad positivity, difference of both, and the results of equivalence tests computed on each electrode in young (top) and older (bottom) participants. The equivalence bounds were computed based on the smallest effect size of interest (SESOI), that is 0.26 in young and 0.29 in older participants. The green color indicates electrodes with a significant equivalence test result, which means that for these electrodes one can reject the hypothesis that the true effect is smaller than d = -SESOI or larger than d = SESOI. In young as well as in older participants, the TOST procedure revealed equivalent centro-parietal topography for P300 and BP.

Note: If the p-value for the equivalence test is less than α = 0.05, then the null hypothesis is rejected meaning that the means are equivalent.

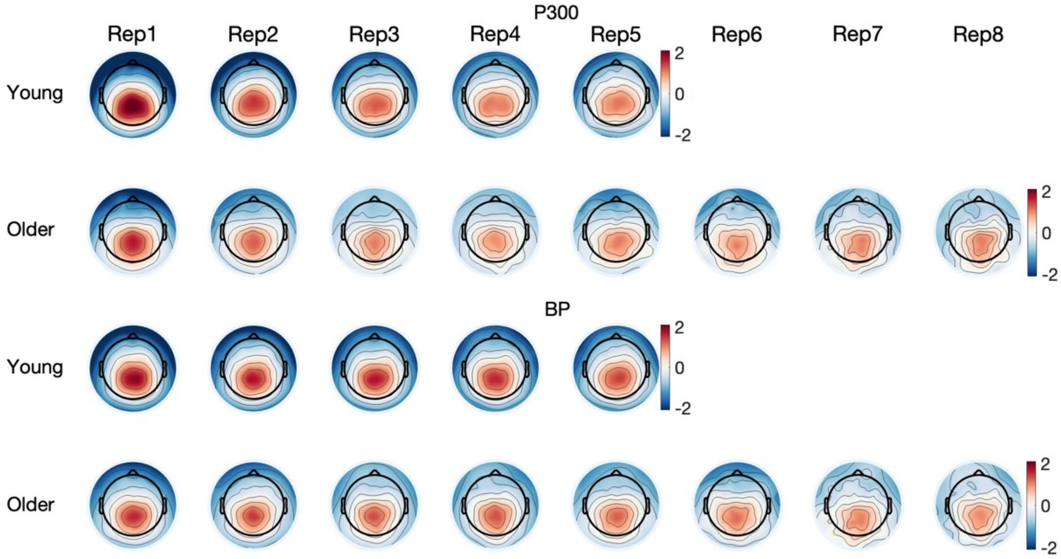


Supplementary Figure 5. Topographical maps of P300 and BP in young and older participants across sequence repetitions. In young, only the first five sequence repetitions were plotted due to low number of stimuli from the sixth sequence repetition on (only for plotting purposes).

**Supplementary Tables (with baseline as predictors)**

Supplementary Table 1. Effects of age group and learning categories on P300 amplitude

| *Variable* | *β* | *SE* | *CI* | *t-value* | *p-value* |
|---|---|---|---|---|---|
| Intercept | 1.13 | 0.10 | 0.93 – 1.33 | 11.23 | 6.5e-13*** |
| Category K | -0.61 | 0.05 | -0.71 – -0.51 | -12.31 | 1.1e-34*** |
| Category UN | -0.11 | 0.06 | -0.23 – 0.02 | -1.66 | 0.096 |
| AgeGroupOld | -0.16 | 0.08 | -0.32 – -0.01 | -2.12 | 0.034* |
| Category K*AgeGroupOld | 0.38 | 0.06 | 0.26 – 0.50 | 6.37 | 2e-10*** |
| Category UN*AgeGroupOld | 0.02 | 0.08 | -0.13 – 0.17 | 0.28 | 0.777 |
| Baseline | -0.29 | 0.01 | -0.32 – -0.26 | -19.51 | 1.9e-84*** |
| Category K*Baseline | 0.08 | 0.02 | 0.05 – 0.11 | 4.62 | 3.8e-06*** |
| Category UN*Baseline | 0.12 | 0.02 | 0.07 – 0.16 | 4.85 | 1.2e-06*** |
| AgeGroupOld*Baseline | 0.16 | 0.02 | 0.12 – 0.20 | 7.93 | 2.3e-15*** |
| Category K * AgeGroupOld * Baseline | -0.13 | 0.02 | -0.17 – -0.08 | -5.36 | 8.5e-08*** |
| Category UN * AgeGroupOld * Baseline | -0.12 | 0.03 | -0.18 – -0.06 | -3.94 | 8.3e-05*** |
| **Variance components** | **SD** | **Goodness of fit** | | | |
| Subject | 0.42 | Log likelihood | | -99919.85 | |
| StimulusNr | 0.15 | | | | |
| RepetitionNr | 0.16 | | | | |
| SequenceNr | 0.05 | | | | |
| Residual | 2.41 | | | | |

*Intercept represents the newly learned (NL) category of young participants.*
**p < 0.05. **p < 0.01. ***p < 0.001.*

Supplementary Table 2. Effects of age group and learning categories on BP amplitude

| ***Variable*** | ***β*** | ***SE*** | ***CI*** | ***t-value*** | ***p-value*** |
|---|---|---|---|---|---|
| Intercept | 0.85 | 0.08 | 0.70 – 1.00 | 11.19 | 2.6e-17*** |
| Category K | -0.45 | 0.05 | -0.54 – -0.35 | -9.30 | 2.6e-20*** |
| Category UN | -0.13 | 0.06 | -0.25 – -0.01 | -2.04 | 0.041* |
| AgeGroupOld | -0.09 | 0.07 | -0.22 – 0.04 | -1.34 | 0.181 |
| Category K*AgeGroupOld | 0.17 | 0.06 | 0.06 – 0.29 | 2.97 | 0.003** |
| Category UN*AgeGroupOld | 0.10 | 0.08 | -0.05 – 0.25 | 1.34 | 0.181 |
| Gender Female | 0.11 | 0.05 | 0.02 – 0.21 | 2.26 | 0.024* |
| Baseline | -0.17 | 0.01 | -0.20 – -0.14 | -11.59 | 5e-31*** |
| Category K*Baseline | -0.00 | 0.02 | -0.04 – 0.03 | -0.21 | 0.832 |
| Category UN*Baseline | 0.05 | 0.02 | 0.00 – 0.10 | 2.05 | 0.04* |
| AgeGroupOld*Baseline | 0.03 | 0.02 | -0.01 – 0.07 | 1.46 | 0.145 |
| Category K * AgeGroupOld * Baseline | 0.01 | 0.02 | -0.04 – 0.05 | 0.28 | 0.776 |
| Category UN * AgeGroupOld * Baseline | -0.07 | 0.03 | -0.13 – -0.01 | -2.37 | 0.018* |
| **Variance components** | **SD** | **Goodness of fit** | | | |
| Subject | 0.33 | Log likelihood | | -99090.4 | |
| StimulusNr | 0.10 | | | | |
| RepetitionNr | 0.08 | | | | |
| Residual | 2.37 | | | | |

*Intercept represents the newly learned (NL) category of young participants.*
*$*p < 0.05$. $**p < 0.01$. $***p < 0.001$.*

Supplementary Table 3. Effects of repetition number and age group on P300 amplitude

| ***Variable*** | ***β*** | ***SE*** | ***CI*** | ***t-value*** | ***p-value*** |
|---|---|---|---|---|---|
| Intercept | 1.37 | 0.07 | 1.24 – 1.51 | 20.39 | 3.1e-44*** |
| RepetitionNr | -0.19 | 0.01 | -0.21 – -0.16 | -13.25 | 1.6e-39*** |
| AgeGroup Old | -0.24 | 0.08 | -0.40 – -0.08 | -2.93 | 0.003** |
| RepetitionNr*AgeGroupOld | 0.11 | 0.02 | 0.08 – 0.14 | 6.74 | 1.8e-11*** |
| Baseline | -0.29 | 0.03 | -0.35 – -0.24 | -10.01 | 2.1e-23*** |
| RepetitionNr*Baseline | 0.02 | 0.001 | 0.01 – 0.04 | 3.21 | 1.3e-03*** |
| **Variance components** | **SD** | **Goodness of fit** | | | |
| Subject | 0.42 | Log likelihood | | -8870.51 | |
| SequenceNr | 0.06 | | | | |
| Residual | 1.03 | | | | |

**p* < 0.05. ***p* < 0.01. ****p* < 0.001.*

Supplementary Table 4. Effects of repetition number and age group on BP amplitude

| *Variable* | *β* | *SE* | *CI* | *t-value* | *p-value* |
|---|---|---|---|---|---|
| Intercept | 0.95 | 0.07 | 0.82 – 1.08 | 14.31 | 1.1e-39*** |
| RepetitionNr | -0.12 | 0.01 | -0.15 – -0.09 | -8.67 | 5.7e-18*** |
| AgeGroup Old | -0.03 | 0.08 | -0.18 – 0.13 | -0.33 | 0.743 |
| RepetitionNr*AgeGroup Old | 0.04 | 0.02 | 0.01 – 0.08 | 2.69 | 0.007** |
| Gender Female | 0.13 | 0.05 | 0.02 – 0.23 | 2.36 | 0.018* |
| Baseline | -0.17 | 0.04 | -0.25 – -0.08 | -3.94 | 8.4e-05*** |
| AgeGroupOld*Baseline | 0.11 | 0.06 | -0.00 – 0.22 | 1.95 | 0.051 |
| RepetitionNr*Baseline | 0.01 | 0.01 | -0.01 – 0.03 | 0.77 | 0.441 |
| AgeGroupOld*RepetitionNr* Baseline | -0.03 | 0.01 | -0.06 – -0.00 | -2.16 | 0.031* |
| **Variance components** | **SD** | **Goodness of fit** | | | |
| Subject | 0.34 | Log likelihood | | -8516.075 | |
| Residual | 0.98 | | | | |

**p* < 0.05. ***p* < 0.01. ****p* < 0.001.*

Supplementary Table 5. Effects of P300 on Knowledge Index

| *Variable* | *β* | *SE* | *CI* | *t-value* | *p-value* |
|---|---|---|---|---|---|
| Intercept | 0.17 | 0.007 | 0.16 – 0.18 | 24.34 | 7e-125*** |
| mP300 | 0.08 | 0.004 | 0.07 – 0.09 | 16.82 | 4.5e-62*** |
| AgeGroup Old | 0.03 | 0.009 | 0.01 – 0.04 | 2.94 | 0.003** |
| mP300*AgeGroup Old | -0.06 | 0.005 | -0.07 – -0.05 | -10.02 | 1.8e-23*** |
| Baseline | -0.01 | 0.006 | -0.02 – 0.00 | -1.72 | 0.085 |
| AgeGroup Old*Baseline | 0.02 | 0.007 | 0.00 – 0.03 | 2.57 | 0.01* |
| **Variance components** | **SD** | **Goodness of fit** | | | |
| Residual | 0.26 | Log likelihood | | -526.10 | |

**p* < 0.05. ***p* < 0.01. ****p* < 0.001.*

Supplementary Table 6. Effects of BP on Learning Index

| *Variable* | *β* | *SE* | *CI* | *t-value* | *p-value* |
|---|---|---|---|---|---|
| Intercept | 0.19 | 0.007 | 0.18 – 0.21 | 28.29 | 1.9e-166*** |
| mBP | 0.06 | 0.005 | 0.05 – 0.07 | 11.13 | 1.7e-28*** |
| AgeGroup Old | 0.004 | 0.009 | -0.01 – 0.02 | 0.4 | 0.688 |
| mBP*AgeGroup Old | -0.04 | 0.007 | -0.06 – -0.03 | 3.81 | 1.4e-04*** |
| Baseline | -0.02 | 0.005 | -0.03 – -0.01 | -3.62 | 2.9e-4*** |
| AgeGroup Old*Baseline | 0.03 | 0.007 | 0.01 – 0.04 | 3.81 | 1.4e-4*** |
| **Variance components** | **SD** | **Goodness of fit** | | | |
| Residual | 0.27 | Log likelihood | | -604.46 | |

**p* < 0.05. ***p* < 0.01. ****p* < 0.001.*
